# Enhanced prediction of breast cancer patient response to chemotherapy by integrating deconvolved expression patterns of immune, stromal and tumor cells

**DOI:** 10.1101/2024.06.14.598770

**Authors:** Saugato Rahman Dhruba, Sahil Sahni, Binbin Wang, Di Wu, Padma Sheila Rajagopal, Yael Schmidt, Eldad D. Shulman, Sanju Sinha, Stephen-John Sammut, Carlos Caldas, Kun Wang, Eytan Ruppin

## Abstract

The tumor microenvironment (TME) is a complex ecosystem of diverse cell types whose interactions govern tumor growth and clinical outcome. While multiple studies have extensively charted the TME’s impact on immunotherapy, its role in chemotherapy response remains less explored. To address this, we developed DECODEM (<u>DE</u>coupling <u>C</u>ell-type-specific <u>O</u>utcomes using <u>DE</u>convolution and <u>M</u>achine learning), a generic computational framework leveraging cellular deconvolution of bulk transcriptomics to associate gene expression of individual cell types in the TME with clinical response. Employing DECODEM to analyze gene expression of breast cancer patients treated with neoadjuvant chemotherapy across three bulk cohorts, we find that the expression of specific immune cells (myeloid, plasmablasts, B-cells) and stromal cells (endothelial, normal epithelial, CAFs) are highly predictive of chemotherapy response, achieving the same prediction levels as the expression of malignant cells. Notably, ensemble models integrating the estimated expression of different cell types perform best and outperform models built on the original tumor bulk expression. These findings and the models’ generalizability are further tested and validated in two single-cell (SC) cohorts of triple negative breast cancer. To investigate the possible role of immune cell-cell interactions (CCIs) in mediating chemotherapy response, we extended DECODEM to DECODEMi to identify such key functionally important CCIs, validated in SC data. Our findings highlight the importance of active pre-treatment immune infiltration for chemotherapy success. DECODEM and DECODEMi are made publicly available to facilitate studying the role of the TME in mediating response in a wide range of cancer indications and treatments.

## INTRODUCTION

The tumor microenvironment (TME) is a complex and multifaceted ecosystem that encompasses a plethora of cell types including immune cells (such as T- and B-lymphocytes, macrophages, dendritic cells etc.), stromal cells (such as fibroblasts, endothelial cells etc.), and the extracellular matrix. It plays crucial roles in regulating tumor progression and response to therapy. Emerging evidence suggest context-specific behavior of the TME as either tumor-suppressive or supportive, thus presenting an attractive target for therapeutic intervention [1][2]. Numerous studies have elucidated the critical role of TME in mediating response to immunotherapy, however, insights into how the TME is relevant to chemotherapy response remains limited. Here we focus on studying the influence of the TME on chemotherapy response in breast cancer (BC), where neoadjuvant chemotherapy (NAC) is used prior to primary surgery to enable tumor downstaging and increase the likelihood of breast-conserving surgery [3][4][5].

Recent studies have underscored the relevance of the BC TME in determining patient response to chemotherapy. For instance, chemotherapy can induce dynamic changes in the TME that lead to an increase in regulatory T-cells and myeloid-derived suppressor-like cells, calling for a balance in immune population for positive treatment outcomes [6]. Lymphocyte density and tumor infiltrating lymphocytes (TILs) have consistently been associated with BC patients achieving pathological complete response (pCR) following NAC and anti-HER2 therapy [7][8][9][10][11]. Additionally, multiple studies have reported the active involvement of TME in therapy resistance [12][13][14], with macrophages, endothelial and BC stem cells promoting chemoresistance [15][16][17], and B-cells and plasma cells displaying varying roles in NAC response [18][19][20]. Based on this evidence, it is apparent that different cells within the TME (and their interactions) contribute significantly to determining chemotherapy response. However, to our knowledge, no study has yet attempted to quantify the contribution and predictive power of each individual cell type within the TME to chemotherapy response in a systematic and rigorous manner. We, therefore, aim to investigate three fundamental questions: (1) Can we apply a rigorous computational approach to delineate the predictive power of cell-type-specific transcriptome in the BC TME from bulk transcriptomics data? (2) Which cell types in the BC TME and the corresponding pathways whose gene expression are most predictive of chemotherapy response? (3) Does aggregating the inferred expression of multiple predictive cell types in the BC TME lead to a more accurate clinical response predictor?

Current advances in precision oncology have highlighted the utility of artificial intelligence [1][2][21][22][23][24] in guiding treatment using diverse data types such as genomics [1][25][26], transcriptomics [29][30][31][32][33], histopathology [34][35][36][37][38][39] or via multimodal approaches [11][27][28][40][41][42]. However, investigations into neoadjuvant therapy response often encounter limitations such as small sample size, treatment heterogeneity, or inadequate capturing of the TME’s complexity, as highlighted by Sammut et al. [11]. In their pioneering work, Sammut et al. presented a novel multiomic predictor of neoadjuvant therapy response in BC [11]. They have leveraged machine learning (ML) to incorporate 34 features extracted from clinical, DNA, RNA, pathology and treatment data to develop a single model to predict patient response across all BC subtypes and different treatment regimens with high accuracy. Expanding on this groundwork and addressing the limitations, our study aims to decouple the cell-type-specific effects to NAC response in the BC TME from transcriptomics alone by focusing on the HER2-negative (HER2-) patients, where NAC is recommended to increase the likelihood of achieving pCR following surgery, especially in triple negative breast cancer (TNBC) [43].

To systematically explore the association between the TME and chemotherapy response, we developed a computational framework, DECODEM (<u>DE</u>coupling <u>C</u>ell-type-specific <u>O</u>utcomes using <u>DE</u>convolution and <u>M</u>achine learning). This framework leverages cellular deconvolution with ML to elucidate the association of the expression of diverse cell types in the TME to patient response to a given treatment. Recognizing that these phenotype effects are intricately regulated by the interactions among relevant cell types within the TME, we further extended DECODEM to DECODEMi, where ‘i’ stands for interaction. DECODEMi incorporates inferred cell-cell interactions (CCIs) to pinpoint the key CCIs influencing treatment response. We demonstrate the utility of our tools in an example scenario of investigating the role of the BC TME in modulating NAC response across multiple bulk and single-cell (SC) cohorts, demonstrating the markedly improved predictive capabilities of cell-type-specific transcriptome over bulk and further pinpointing the notable CCIs. A recent study by Kester et al. investigated the association of cellular compositions of the TME with patient survival in BC [44], but to our knowledge, the current study is the first to analyze the cell-type-specific transcriptomic profiles in the BC TME to objectively quantify their association with treatment response. Together our findings point to the necessity of an active pre-treatment immune system to facilitate positive treatment outcome, as supported by Sammut et al. [11]. From a methodological perspective, our study provides tools to systematically assess the role of individual cellular components within the TME in anticancer therapy response (DECODEM) and to identify the prominent cell-to-cell communications associated with this response (DECODEMi).

## RESULTS

### Overview of the analysis

DECODEM and DECODEMi are new computational frameworks designed to explore the predictive powers of individual cell types and relevant cell-cell interactions, respectively, within the TME to response to therapy. They harness the prowess of CODEFACS and LIRICS, two recent tools from our laboratory [45]. CODEFACS facilitates the extraction of cell-type-specific expression profiles for each patient from their bulk gene expression, whereas LIRICS discerns the activity of cell-type-specific ligand-receptor interactions within their TME. In essence, DECODEM takes pre-treatment bulk gene expression data of a patient tumor sample as the input. It then uses deconvolution and machine learning to generate a score for each cell type indicating the likelihood that the patient will respond to the therapy. The predictive performance of these scores are quantified by using the standard classification metrics i.e., area under the receiver operating characteristics curve (AUC), average precision (AP; equivalent to area under the precision-recall curve) and diagnostic odds ratio (DOR) metrics. DECODEMi then follows through to associate this response with CCIs in the TME to extract the most predictive interactions among these cell types.

Specifically, DECODEM comprises of two sequential steps (Figure 1D): First, for each bulk expression cohort, it generates nine cell-type-specific expression profiles for each sample across the whole transcriptome using CODEFACS with a BC-specific signature generated using scRNA-seq data (n = 26) from Wu et al. [46] (see METHODS) with annotations provided for nine major cell types: B-cells, cancer-associated fibroblasts (CAFs), cancer epithelial (malignant), endothelial, myeloid, normal epithelial, plasmablasts, perivascular-like cells (PVL) and T-cells; Second, by leveraging these deconvolved expression profiles, it generates nine cell-type-specific predictors of clinical response (classifying responders vs. non-responders e.g., pCR vs. residual disease) using a multi-stage ML pipeline. Within this pipeline, we performed rigorous feature selection to keep the top ‘m’ genes (2 ≤ m ≤ 25; the upper limit is chosen to avoid overfitting by considering the confidently inferred genes) per cell type before feeding their expression into an unweighted ensemble classifier consisting of four algorithms: regularized logistic regression, random forest, support vector machine and XGBoost. The pipeline was trained with repeated three-fold cross-validation (CV) to optimize model hyperparameters. Additionally, we built a baseline predictor with bulk expression using the same ML pipeline to serve as a control, comparative rod (we also built a second baseline predictor with the estimated cell fractions that consistently underperformed the bulk baseline, thus opting to reporting only the latter). Finally, as a sanity check, we performed a benchmark analysis for CODEFACS deconvolution accuracy in breast cancer, our application domain. We used five BC benchmark cohorts where patient pseudobulk samples were generated via mixing individual cells from Wu et al. [46] (see METHODS). CODEFACS was then applied to these simulated bulk expression profiles with known ground truth. The results reassuringly showed that CODEFACS achieved high accuracy and successfully inferred thousands of genes accurately across cell types, thereby ensuring the robustness of the deconvolution (Supplementary Figure 1).

**Figure 1:**
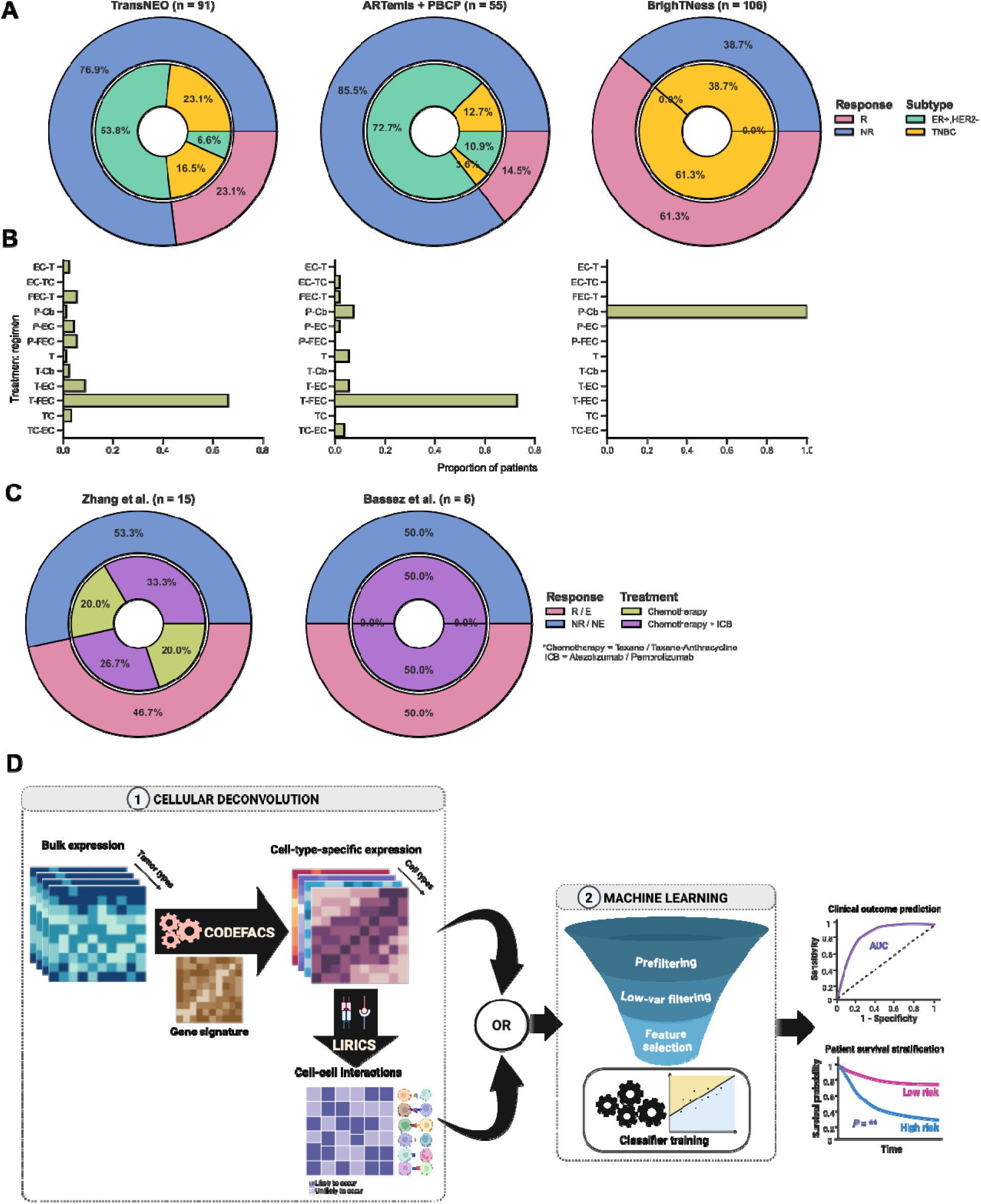
Overview of the methodology: cohorts and analysis. **A-B.** Overview of the bulk cohorts used for analysis i.e., the ratio of responders to non-responders (R:NR), clinical subtype distribution (**A**) and available treatment regimens (B) acros TransNEO, ARTemis + PBCP and BrighTNess. TransNEO was used for model development and the latter two were used for validation. ER+,HER2- and TNBC refer to the ER-positive, HER2-negative and triple negative breast cancer subtypes, respectively. C, Cb, E, F, P and T stand for cyclophosphamide, carboplatin, epirubicin, 5-flurouracil, paclitaxel and docetaxel (taxotere), respectively. **C**. Overview of the single-cell (SC) TNBC cohorts used for external validation. The patients in Zhang et al. were analyzed separately based on treatment. For Bassez et al., T-cell clonotype expansion (E) and non-expansion (NE) was used as a surrogate for patient response. ICB stands for immune checkpoint blockade (anti-PD1 / anti-PD-L1) therapy. **D**. The analysis pipeline, **DECODEM** and **DECODEMi**. First, we apply CODEFACS on bulk expression to infer cell-type-specific expression profiles or LIRICS for the cell-cell interactions (CCIs) present in the tumor microenvironment. Second, we train a multi-stage ML pipeline with these cell-type-specific expression profiles to build cell-type-specific clinical response predictors (DECODEM) or the binary CCI profile to build a CCI-based predictor (DECODEMi). For validation, we deploy the first stage above (bulk) or a customized pipeline (SC) to prepare cell-type-specific expression or CCIs and input to the trained predictors to evaluate model in terms of the area under the receiver operating characteristics curve (AUC) or stratify patient survival using Kaplan-Meier analysis.

To assess the performance of DECODEM and evaluate the contribution of each cell type, we analyzed bulk transcriptomics data from three BC cohorts with different ratios of responders to non-responders (R:NR) to NAC (Figure 1A) in two scenarios: (a) model development and cross-validation with TransNEO (n = 94, R:NR = 22:72), recently published by Sammut et al. [11] and (b) external validation on ARTemis + PBCP (n = 55, R:NR = 8:47) [47] and BrighTNess (n = 106, R:NR = 65:41) [48]. TransNEO and ARTemis + PBCP covered all major BC subtypes among which we focused on the HER2-patients, treated with various chemotherapy regimens [11][47] (Figure 1A-B), whereas BrighTNess included only TNBC patients from the arm B of the trial, treated with paclitaxel and carboplatin with a placebo for veliparib [48]. We further validated DECODEM’s findings in two independent TNBC single-cell cohorts (Figure 1C): Zhang et al., which consisted of 15 patients receiving either paclitaxel alone (R:NR = 3:3) or in combination with the anti-PD-L1 agent, atezolizumab (R:NR = 4:5) [49] and Bassez et al., which consisted of six patients (R:NR = 3:3; taking T-cell clonotype expansion as a surrogate for patient response) receiving a taxane – anthracycline based NAC before the anti-PD1 agent, pembrolizumab [50]. Notably, we took the Sammut et al. predictor built with 18 extracted features from clinical and RNA-seq data (e.g., not including the DNA-seq data) as a comparative rod that achieved an AUC of 0.88 in CV and 0.89 in validation with ARTemis + PBCP [11], outperforming a baseline predictor (AUC = 0.7) built with seven clinical features including ER and HER2 statuses, which are routinely used to guide clinical decisions [43].

To investigate the impact of incorporating the expression profiles of multiple predictive cell types together, as indicated by our third question, we applied two extensions to DECODEM: (i) append the cell-type-specific expression profiles for two and three most predictive cell types together to train a multi-cell-ensemble ML pipeline and (ii) extend to DECODEMi to incorporate the inferred CCIs to train a CCI-based ML pipeline (Figure 1D). The former model maximizes the predictive performance by leveraging complementary information from different cell types while the latter identifies the most predictive cell-to-cell communications for clinical response, thereby providing deeper insights to the underlying mechanism.

### Gene expression of immune and stromal cells predicts response to neoadjuvant chemotherapy as accurately as the expression of malignant cells

To assess the impact of individual cell types on chemotherapy response, we applied DECODEM to TransNEO in a CV analysis. Our analysis demonstrated significant differences (one-tailed Wilcoxon rank-sum test P-value ≤ 0.05) in the prediction scores between responders and non-responders across 91 patients (i.e., patients with available prediction scores from Sammut et al.) in TransNEO (Figure 2A), with seven cell types exhibiting enhanced capabilities of identifying responders over the bulk and Sammut et al. predictors (Figure 2D). Notably, the immune cells (myeloid, plasmablasts, B-cells) contribute as prominently as malignant (cancer epithelial) and stromal cells (endothelial, normal epithelial, CAFs; Figure 2D). Our subsequent analyses further showed that these seven ‘prominent’ cell types accurately identify true responders to chemotherapy with high average precision (Figure 2G) and diagnostic odds ratios (DOR; Figure 2J). We also investigated if training strategy impacts model performance by using a leave-one-out CV instead of three-fold CV for hyperparameter tuning (maximizing accuracy) and obtained similar performance (Supplementary Figure 2), supporting the robustness of our chosen strategy.

**Figure 2:**
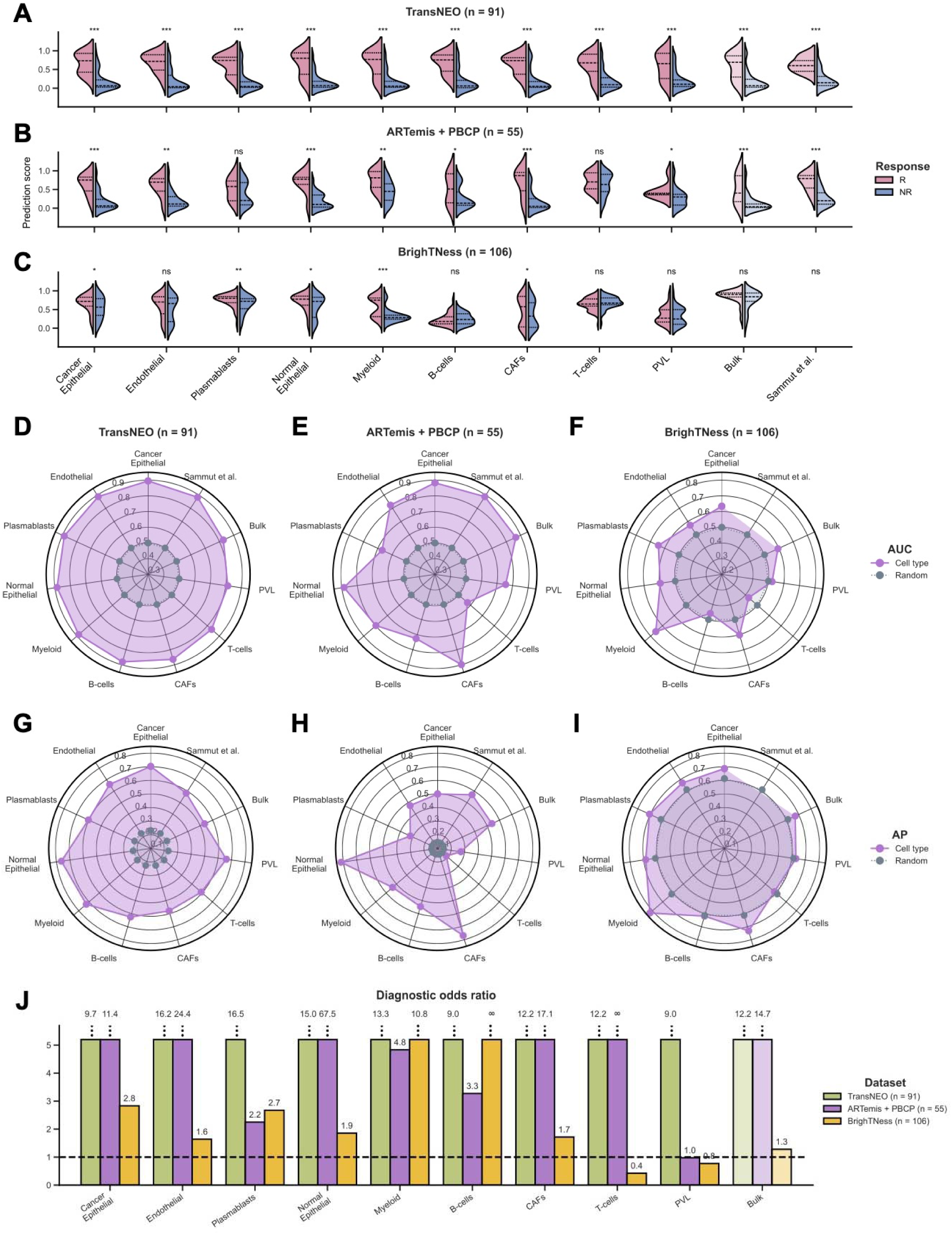
The prominent cell types mediating chemotherapy response in breast cancer tumor microenvironment. **A-C**. Comparison of prediction scores for nine cell-type-specific, bulk and Sammut et al. (for first two cohorts) predictors across TransNEO (A), ARTemis + PBCP (B) and BrighTNess (C). R and NR stand for responders and non-responders, respectively. The differences between the prediction scores were computed by using a one-tailed Wilcoxon rank-sum test (***, **, * and ‘ns’ denote P ≤ 0.001, P ≤ 0.01, P ≤ 0.05 and P > 0.05, respectively). Cell types are ranked by their AUC values for TransNEO in a descending order. **D-I.** Comparison of model performance for nine cell-type-specific, bulk and Sammut et al. (for first two cohorts) predictors across TransNEO (D,G), ARTemis + PBCP (E,H) and BrighTNess (F,I). AUC and AP stand for the area under the receiver operating characteristics curve and average precision, respectively. ‘Random’ denotes a random predictor (AUC = 0.5, AP = fraction of responders). Cell types are ranked by their AUC values for TransNEO in a descending order in the counterclockwise direction. **J**. Comparison of diagnostic odds ratios (DOR) for nine cell-type-specific and bulk predictors across the three cohorts. The dotted line represents a random predictor (DOR = 1.0). Cell types are ranked by their AUC values in TransNEO a descending order.

These cell types have frequently been reported to influence clinical response in BC, particularly in the context of chemotherapy-induced immunogenic cell death as the proposed mechanism [52][11]. Changes in the immune population, such as an increase in regulatory T-cells and myeloid-derived suppressor-like cells, and a decrease in CD8 T-cells, have been associated with achieving pCR following NAC [6]. Myeloid cells have been implicated in therapy response through interactions with the malignant and endothelial cells, inducing chemoresistance via the CXCL1/2 – S100A8/9 loop, whereas tumor-associated macrophages have been linked to adverse outcomes and chemoresistance through survival factor secretion, anti-apoptotic pathway activation and modulation of signaling pathways [6][15][16]. VEGF-mediated upregulation of survivin in endothelial cells was further implicated in playing central roles in chemoresistance [15]. Additionally, plasmablasts, as precursors of plasma cells, have been associated with better prognosis [19][20], while B-cells have shown both pro- and anti-tumor roles [18][19][20].

We further tested and validated the predictive power of these cell types in external validation analyzing the ARTemis + PBCP (n = 55) and BrighTNess (n = 106) cohorts, observing fairly consistent results (Figure 2B-C, E-F, H-J). The differences in score distributions across cohorts (Figure 2A-C) likely stem from both biological variations due to cohort differences in tumor subtypes, stages, tumor sample cell composition and ER / HER2 status, and variations arising from out-of-distribution generalization effects (e.g., different R:NR ratios across cohorts), a well-known challenge in ML modeling. In the ARTemis + PBCP cohort (dominated by ER-positive, HER2-negative (ER+,HER2-) BC), the prominent contributors to chemotherapy response were CAFs, normal epithelial and malignant cells (Figure 2E,H), whereas in the TNBC-specific BrighTNess cohort, the key players were myeloid, plasmablasts and malignant cells (Figure 2F,I). For both cohorts, our prominent cell types accurately identified the responders with notably high (>1.6) DOR values (Figure 2J) with variations observed across cohorts due to the effect of cohort composition (i.e., ARTemis + PBCP has few responders with R:NR = 8:47, as opposed to BrighTNess with R:NR = 65:41). When stratifying TransNEO and ARTemis + PBCP cohorts by the available BC subtypes, our cell-type-specific models retain good performance across subtypes (Supplementary Figure 3A-D), although the models for ER+,HER2-BC were more predictive than TNBC, possibly due to sample size limitations and the increased heterogeneity in TNBC. These findings support the prominent involvement of immune and stromal cells with the malignant cells in mediating patient response to chemotherapy within the TME, with variations observed likely due to tumor subtype composition, stage and ER / HER2 status differences across cohorts.

Our analysis did not identify T-cells as a prominent predictive cell type, as their expression provide marginal to no enhancement in predictive power over bulk expression (Figure 2D-J). This lack of signal may stem from aggregating different T-cell subtypes together (due to limits on the resolution of the deconvolution, which breaks down for low-abundance cell types). To study this further, we performed an enrichment analysis to delineate the contributions of CD4 and CD8 T-cells to chemotherapy response. However, the expression of each of these cell types still exhibited limited stratification power to distinguish responders from non-responders across the three bulk cohorts (Supplementary Figure 4A-D). We further complemented this expression-based analysis with an association analysis of cell abundance and clinical response, revealing that the overall T-cell abundance exhibited predictive power similar to or even higher than CD4 and CD8 subtypes (Supplementary Figure 4E-F), albeit still notably lower than that of bulk.

### An ensemble model incorporating the expression of immune and stromal cells significantly boosts the predictive power of patient response

To further enhance DECODEM’s performance, we aggregated multiple individual cell types together by appending their expression profiles as input features. We explored 35 combinations of two to three cell types exhaustively, assessing whether a selective aggregation of multi-cell-type gene expression yields better predictive scores than the single-most predictive cell type. This comprehensive analysis achieved a boost in predictive performance for both ARTemis + PBCP and BrighTNess, with significant overlaps observed in top multi-cell-type ensembles across cohorts (Figure 3A-F). Remarkably, the ensemble of immune and stromal cells i.e., the aggregation of the expressions of endothelial, myeloid and plasmablasts exhibited the strongest capability in identifying chemotherapy responders (ARTemis + PBCP: AUC = 0.94; BrighTNess: AUC = 0.75; Figure 3D). Additionally, the immune-cell-ensemble comprising myeloid and plasmablasts displayed high stratification (ARTemis + PBCP: AUC = 0.92; BrighTNess: AUC = 0.73; Figure 3A). As before, these multi-cell-ensembles also yielded high average precision (Figure 3B,E) and notably high (>5.0) DOR values (Figure 3C,F). These findings testify to the complementary contributions of diverse cell types within the TME to chemotherapy response. However, the top ensemble for BrighTNess provided a performance similar to the top cell type, myeloid (AUC = 0.76; Figure 2F), suggesting that the interactions among cell types are less prominent in this cohort compared to the others, expanded further in Figure 4A-B. The subtype-specific stratification of ARTemis + PBCP reinforced the superior predictive capabilities of these ensembles across two subtypes while recapitulating their diminished impact in TNBC (Supplementary Figure 3E-F).

**Figure 3:**
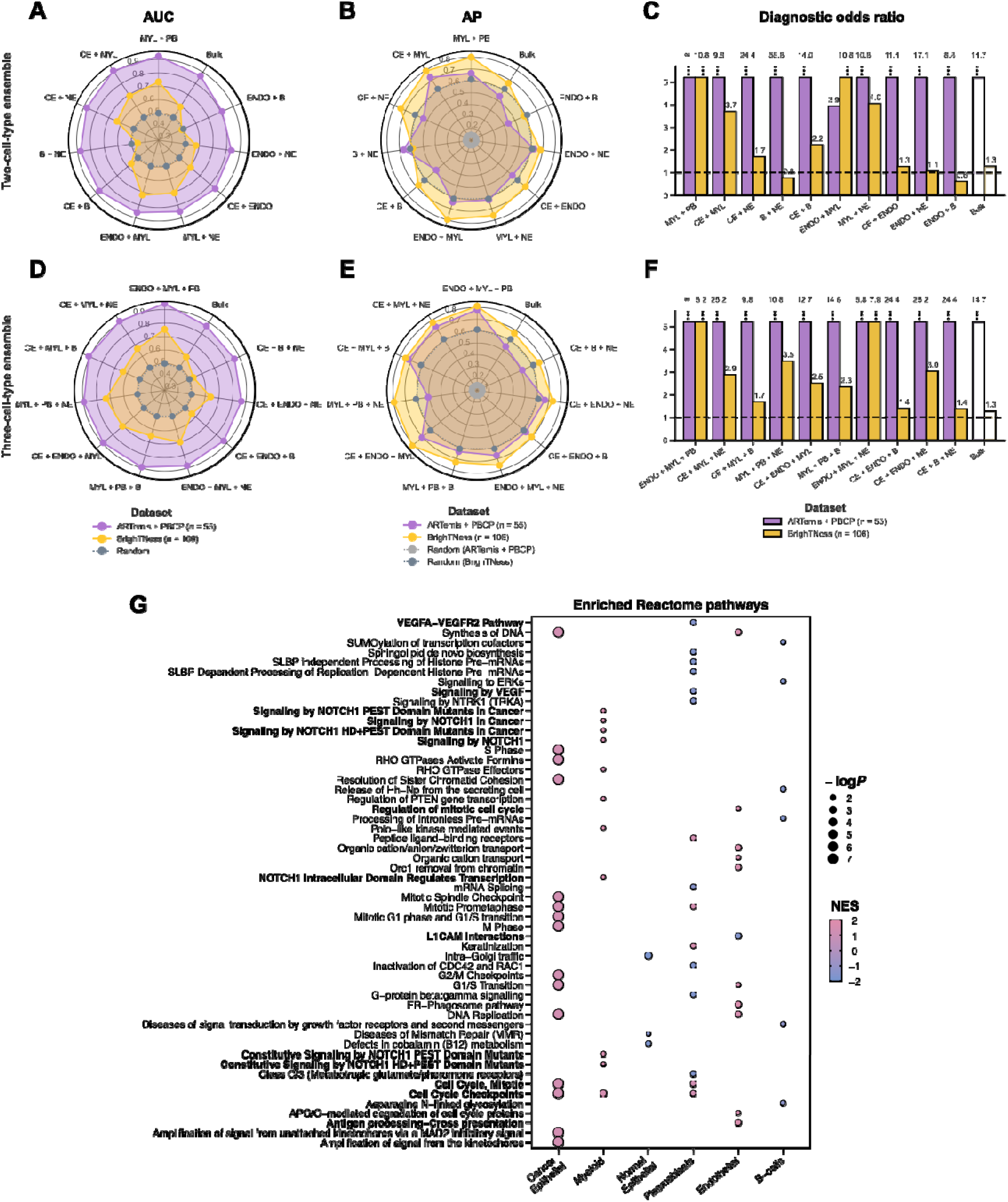
The prominent multi-cell-type ensembles mediating chemotherapy response in breast cancer tumor microenvironment. **A-C.** Comparison of external validation model performance for the 10 most predictive ensembles comprising of two cell types with the bulk predictor. B, CE, ENDO, MYL, NE and PB stand for B-cells, cancer epithelial (malignant), endothelial, myeloid, normal epithelial and plasmablasts, respectively. AUC and AP stand for the area under the receiver operating characteristics curve and average precision, respectively. ‘Random’ denotes a random predictor (AUC = 0.5, AP = fraction of responders). The dotted line in C represents the diagnostic odds ratio (DOR) for a random predictor (DOR = 1.0). Ensembles are ranked by their mean AUC values across cohorts in a descending order in the counterclockwise direction. **D-F.** Comparison of external validation model performance for the 10 most predictive ensembles comprising of three cell types with the bulk predictor. Ensembles are ranked by their mean AUC values across cohorts in a descending order in the counterclockwise direction. **G**. The significantly enriched Reactome pathways (FDR-adjusted P ≤ 0.2) across six prominent cell types. For each cell type, only the 10 most relevant pathways are displayed. NES and logP stand for the normalized enrichment score from GSEA and log-scaled P-value, respectively.

**Figure 4:**
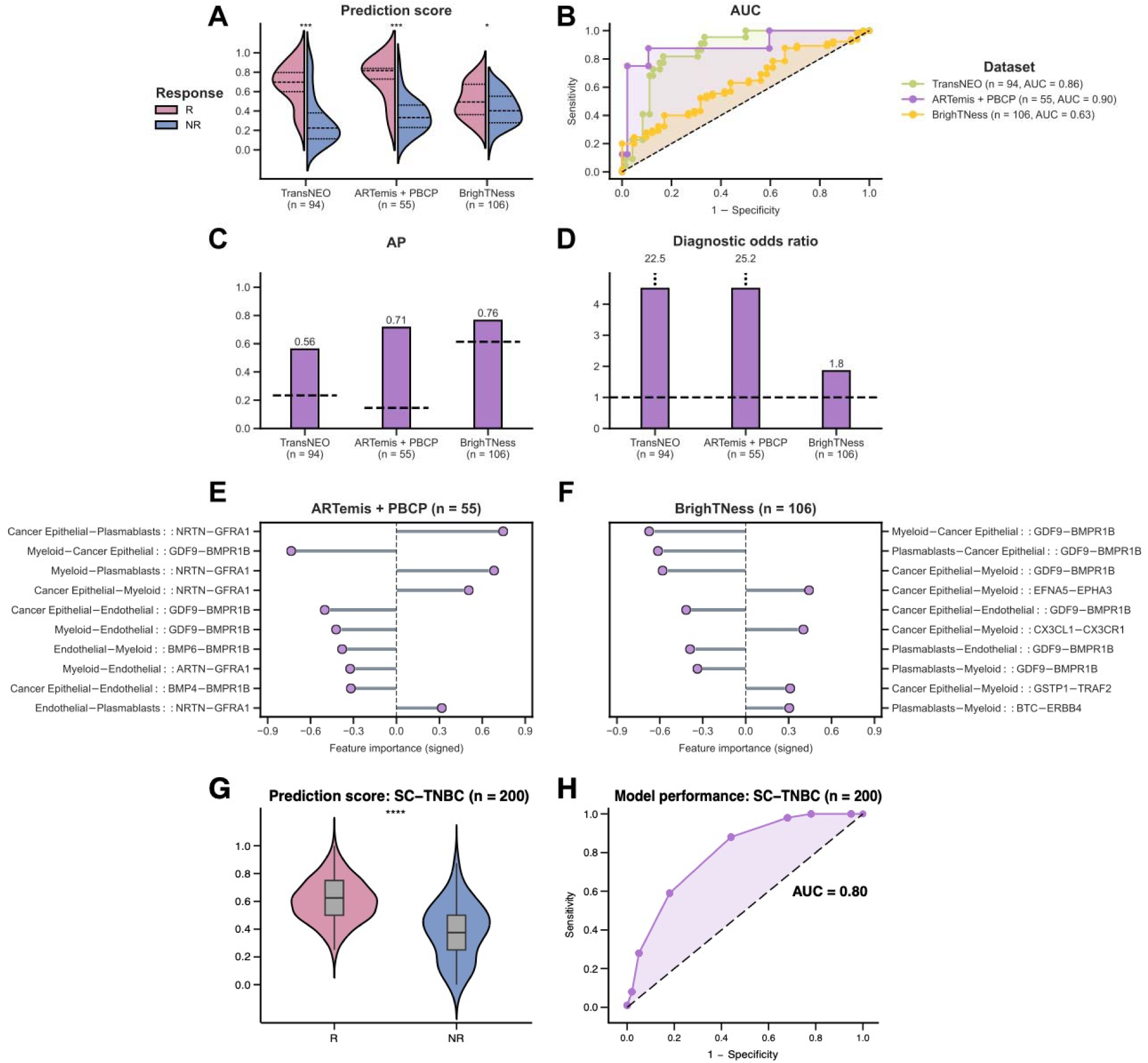
The prominent cell-cell interactions (CCIs) mediating chemotherapy response in breast cancer tumor microenvironment. **A-D.** Comparison of prediction scores (A) and model performance (B-D) for DECODEMi CCI-based predictor across TransNEO, ARTemis + PBCP and BrighTNess. R and NR stand for responders and non-responders, respectively. The differences between the prediction scores were computed by using a one-tailed Wilcoxon rank-sum test (****, *** and * denote P ≤ 0.0001, P ≤ 0.001 and P ≤ 0.05, respectively). AUC and AP stand for the area under the receiver operating characteristics curve and average precision, respectively. The dotted lines in C-D represent the AP and diagnostic odds ratio (DOR) values for a random predictor (AP = fraction of responders, DOR = 1.0). **E-F.** The 10 most predictive CCIs in external validation with ARTemis + PBCP (E) and BrighTNess (**F**). Each CCI is displayed as a quadruplet where the first pair containing the ligand and receptor cell types (separated by ‘–’) is separated by ‘::’ from the second pair containing the corresponding ligand and receptor genes (separated by ‘–’). Feature importance values were computed by using Gini impurity (the built-in importance measure in random forest where a higher value indicates a higher importance and vice versa). Feature directionalities were computed by using a two-sided Fisher’s enrichment analysis (i.e., positive when odds ratio > 1 and vice versa). **G-H**. Prediction scores (G) and model performance (H) in external validation with SC-TNBC (n = 200) using the top 170 response-relevant CCIs among B-cells, myeloid and T-cells, as identified by DECODEMi in bulk.

We conducted gene set enrichment analysis (GSEA) with Reactome pathways [53] to explore the functional contributions of these prominent cell types in chemotherapy response prediction. This analysis revealed the enrichment of cell cycle checkpoint and DNA replication pathways across different cell types (false discovery rate (FDR)-adjusted P-value ≤ 0.2; Figure 3G) as the most common contribution. Dysregulation of these pathways are frequent in BC and promotes tumor proliferation [54][55][56], making them valuable prognostic markers and therapeutic targets [57][58][59][60]. Immune cells exhibited enrichment in signal transduction pathways (e.g., VEGFA – VEGFR2 and VEGF signaling pathways in plasmablasts and multiple NOTCH1 signaling pathways in myeloid), implicated in malignant progression and therapy resistance, particularly in TNBC [61][62][63][64][65][66][67]. Stromal endothelial cells showed enrichment in development- and immune-related pathways (e.g., L1CAM interactions and antigen processing & presentation), contributing to aggressive BC progression and immune evasion with implications for both chemo and immunotherapy [68][69][70][71][72][73][74].

We next assessed DECODEM’s performance at a higher resolution considering the prevalent chemotherapy regimen in each cohort. TransNEO and ARTemis + PBCP predominantly employed T-FEC (n = 61 and 40, respectively; combining docetaxel (taxotere), 5-flurouracil, epirubicin and cyclophosphamide), while BrighTNess used a regimen of paclitaxel and carboplatin (Figure 1B). All six drugs display immunogenic effects by stimulating anticancer immunity via either on-target (releasing immunostimulatory molecules from malignant cells) or off-target (promoting activation of effector cells and diminishing immunosuppressive cells) effects [74]. Remarkably, across all three cohorts spanning two BC subtypes and two treatment regimens, DECODEM consistently identified myeloid, plasmablasts and B-cells to stratify responders to immunogenic drug regimens, alongside the malignant cells (Supplementary Figure 5A-C). For ARTemis + PBCP (dominated by ER+,HER2-BC), although the bulk mixture displayed comparable or superior predictive power than each of these individual cell types, their ensembles markedly boosted the performance, supporting the presence of prominent CCIs involving these cell types that may be associated with the likelihood of response (Supplementary Figure 5D). In addition, the ensemble of endothelial, myeloid and plasmablasts retains its highest performance, further underscoring the critical roles of stromal – immune interactions. For BrighTNess, the individual immune cell types showed notably strong predictive abilities, with less pronounced CCI effects (Supplementary Figure 5E, Figure 4B-D). Collectively, these findings corroborate the immunogenic cell death effects mediated by these chemotherapies [74], reinforcing previous suggestions by Sammut et al. [11].

### Identifying key cell-cell interactions associated with chemotherapy response

To investigate the impact of the presence of interactions among different cell types in modulating chemotherapy response, we use DECODEMi to analyze CCIs (inferred using LIRICS [45], starting with 2,422 ligand – receptor (L-R) pairs from Ramilowski et al. [51]; see METHODS) that are predictive of patient response. As before, we conducted two analyses: (a) model development and cross-validation with TransNEO (n = 94) and (b) external validation on ARTemis + PBCP (n = 55) and BrighTNess (n = 106). As a preliminary analysis, we found that the CCIs encompassing the four most prominent cell types (cancer epithelial, endothelial, myeloid and plasmablasts) alone exhibited a classification power (one-tailed Wilcoxon rank-sum test P-value ≤ 0.05) comparable to all available CCIs across the three cohorts, further testifying to the influential roles of these prominent cell types in patient response (Figure 4A-D, Supplementary Figure 6A-D). The CCIs involving these four cell types markedly improved the predictive power of CCI-based predictors over bulk in both TransNEO and ARTemis + PBCP (Figure 4B-D). However, the CCI performance was generally lower than the top predictive myeloid cells in BrighTNess (Figure 4B-D), which can again be attributed to cohort composition, stage and treatment-specific variations. It is worth noting that BrighTNess included only later-stage TNBC patients treated with paclitaxel and carboplatin, a combination for which very few samples were available in training (Figure 1A-B). These results reinforced that leveraging information from multiple cell types can notably improve clinical response prediction and the intercellular crosstalk may be involved in mediating chemotherapy response within the TME.

To further assess the generalizability of DECODEMi, we evaluated its effectiveness on an external single-cell TNBC cohort (SC-TNBC; sourced from Zhang et al. [48]). To overcome small sample size limitation, we generated 200 ‘pseudopatients’ treated with NAC (R:NR = 100:100) by downsampling the 28,209 cells across the six TNBC patients from Zhang et al. who were treated with paclitaxel [49]. Focusing on the CCIs involving the three cell types present in both SC and deconvolved bulk data (B-cells, myeloid and T-cells), we identified the 170 most predictive CCIs (covering 134 L-R pairs) from TransNEO and computed prediction scores for the SC-TNBC cohort by aggregating the activated CCIs for each pseudopatient (as inferred by SOCIAL [75]; see METHODS). Our analysis highlighted the significant predictive capabilities of these top CCIs (one-tailed Wilcoxon rank-sum test P-value ≤ 0.001; AUC = 0.8; Figure 4G-H), affirming DECODEMi’s ability to identify the CCIs predictive of chemotherapy response from bulk expression.

By computing feature importance from DECODEMi, we identified the most predictive cell-to-cell communications across ARTemis + PBCP and BrighTNess, finding a significant overlap in top CCIs (Figure 4E-F). Remarkably, the interaction between ‘GDF9’ (growth differentiation factor 9) and ‘BMPR1B’ (bone morphogenetic protein receptor 1B), both members of the TGF-β superfamily (the corresponding functional protein-protein interaction (PPI) network, obtained from STRING [76], is provided in Supplementary Figure 6E) with roles in reproductive biology and signal transduction [77][78][79][80], emerged as a key CCI with the same directionality across multiple cell types. These genes have been implicated in BC prognosis and therapeutic outcomes [81][82][83], with BMPR1B exhibiting dual roles in chemotherapy response [84][85][86]. Another top CCI involved the renowned GDNF – RET signaling pathway interactions between ‘NRTN’ (neurturin) or its paralog ’ARTN’ (artemin) and ‘GFRA1’ (GDNF family receptor alpha 1) in multiple cell types (Figure 4E-F; see PPI network in Supplementary Figure 6F), which are all members of the glial cell line-derived neurotrophic factor (GDNF) family within the TGF-β superfamily with roles in neuronal survival, growth, and differentiation [87]. The GDNF family exhibits diverse roles in cancer [88][89], including an association with therapy resistance in ER+ BC, highlighting GFRA1 as a potential target [90][91][92][93]. Additionally, the well-studied chemokine interaction between ‘CX3CL1’ (C-X3-C chemokine ligand 1) and ‘CX3CR1’ (C-X3-C chemokine receptor 1) [94] is prominent in BrighTNess (Figure 4F; see PPI network in Supplementary Figure 6G) and known to be involved in progression and metastasis across multiple cancers including BC, presenting a promising pan-cancer target [95][96][97][98][99][100].

Overall, DECODEMi application results in: (i) a simpler predictor leveraging CCIs that improves patient stratification compared to bulk transcriptomics and (ii) the identification of CCIs predictive of treatment response, whose activation is associated with pro-cancerous roles, thus offering potential novel therapeutic targets.

### DECODEM models predict patient response to chemotherapy and ICB therapy as well as T-cell clonotype expansion from single-cell transcriptomics

To further assess the generalizability of DECODEM, we evaluated the predictive capabilities of the cell-type-specific models with single-cell transcriptomics. Due to the scarcity of publicly available SC cohorts with NAC response, we extended our analysis to predict response to both neoadjuvant chemotherapy and immune checkpoint blockade (ICB) therapy, shown to improve pCR and approved recently as a first-line treatment in early-stage TNBC [101][102][103].

We first analyzed the Zhang et al. cohort [49] containing the expression of B-cells, myeloid and T-cells for 15 TNBC patients, either treated with paclitaxel alone (n = 6) or combined with the anti-PD-L1 agent, atezolizumab (n = 9). We stratified patients by treatment and applied our established DECODEM models for these three cell types to predict response. Notably, these models were trained on deconvolved TransNEO data without any further training on SC data. Our findings revealed that for both intervention types, pre-treatment expression of B-cells exhibited the highest responder stratification capability (one-tailed Wilcoxon rank-sum test P-value ≤ 0.05; Figure 5A; Supplementary Figure 7A-B) and predictive potential (Figure 5B, Supplementary Figure 7D-E). This is consistent with the findings from the original study that B-cells at baseline display the highest association with favorable response to both chemotherapy and ICB therapy [49]. Further, our analysis displayed a moderate predictive power for T-cells (AUC: chemotherapy = 0.67; chemotherapy + ICB = 0.67; Supplementary Figure 7D-E), the major hit from Zhang et al. [49], despite a weak T-cells-specific predictor (Figure 2D-F).

**Figure 5:**
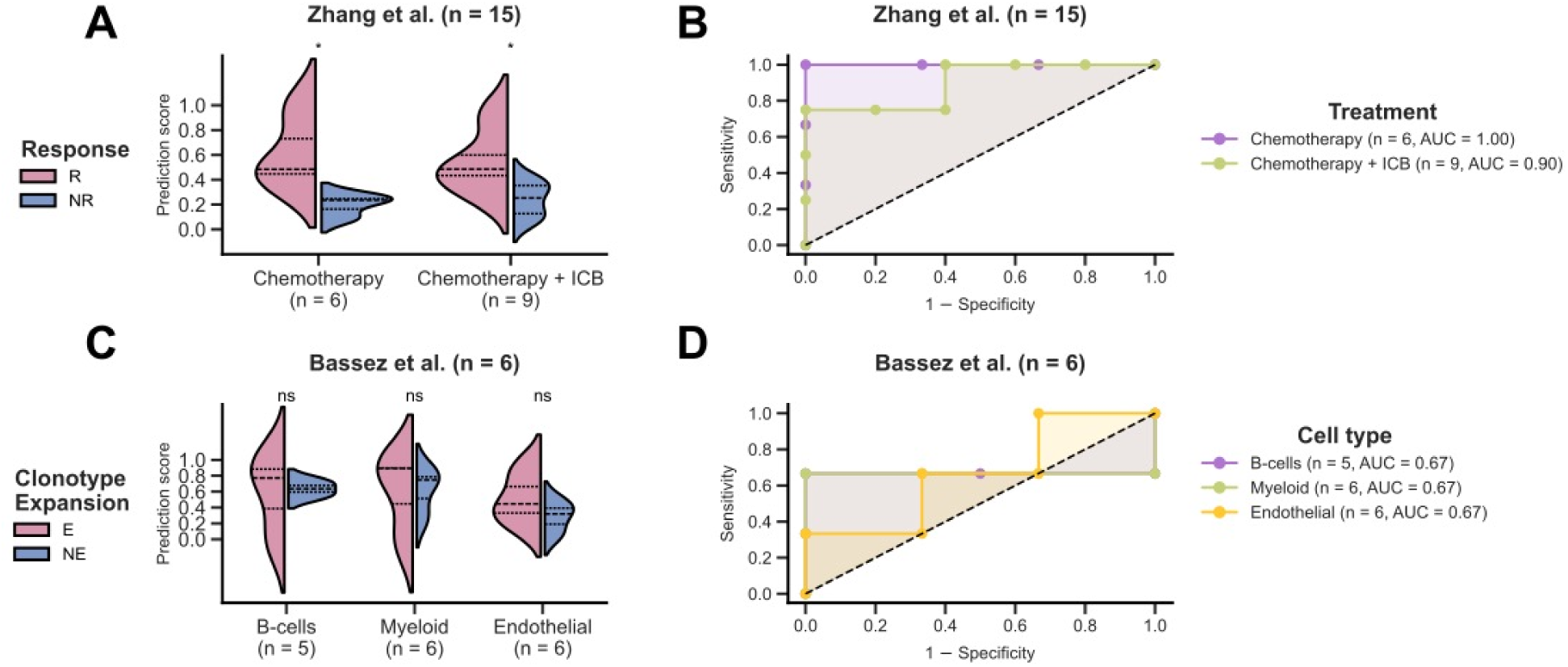
DECODEM generalizes to single-cell (SC) for clinical response prediction and for patient survival stratification. **A-B.** Prediction scores (A) and model performance (B) for B-cells in external validation with Zhang et al. SC cohort for neoadjuvant chemotherapy and immune checkpoint blockade (ICB) therapy response prediction. T-cell clonotype expansion is taken as a surrogate for response in Bassez et al. R and NR stand for responders and non-responders, respectively. The differences between the prediction scores were computed by using a one-tailed Wilcoxon rank-sum test (* and ‘ns’ denote P ≤ 0.05 and P > 0.05). AUC stands for the area under the receiver operating characteristics curve. **C-D.** Prediction scores (C) and model performance (D) across cell types in external validation with Bassez et al. SC cohort for T-cell clonotype expansion prediction post-treatment with neoadjuvant chemotherapy and ICB. E and NE stand for expanders and non-expanders, respectively.

We next analyzed the Bassez et al. cohort [50] containing the expression of B-cells, CAFs, malignant, endothelial, myeloid and T-cells for six TNBC patients, treated with chemotherapy (taxane – anthracycline based) and the anti-PD1 agent, pembrolizumab. This study measured T-cell clonotype expansion using scTCR-seq to identify patients with post-treatment clonotype expansion vs. non-expansion (E vs. NE), which we take as a surrogate for response. Applying the established cell-type-specific models, our analysis revealed myeloid, B-cells and endothelial as the most predictive cell types at baseline (Figure 5C-D, Supplementary Figure 7C,F), consistent with the original findings that certain pre-treatment macrophages are associated with T-cell expansion while B-cells in tertiary lymphoid structures can impact ICB response [50]. Notably, the study pointed to half of the NE patients having higher percentages of baseline T-cells (that failed to expand post-ICB) [50], likely contributing to our finding of limited T-cells predictive power.

These findings attest to the generalizability and robustness of DECODEM, highlighting its potential in capturing the treatment-invariant properties of the BC TME to predict response to the combination of chemotherapy and ICB therapy through SC transcriptomics.

Similarly, we explored the prognostic capabilities of the DECODEM scores by using the early-stage HER2-breast cancer patients in TCGA-BRCA (n = 484) and achieved the most effective survival stratification across the malignant cells for both overall survival (Supplementary Figure 8) and progression-free interval (Supplementary Figure 9). These results attest that the cell-type-specific models can indeed capture the treatment-invariant properties of BC TME to indicate patient survival despite not explicitly training for it.

## DISCUSSION

Our study introduces DECODEM, a transcriptomics-based modeling framework harnessing CODEFACS, a cutting-edge deconvolution tool, with a multi-stage machine learning pipeline to systematically assess the cell-type-specific contributions to the prediction of clinical response to treatment. As a representative scenario, we apply it to build predictors of patient response to neoadjuvant chemotherapy in breast cancer. We further present DECODEMi, a companion tool that identifies the key cell-cell interactions associated with such clinical response. A recent study by Kester et al. investigated the association of cellular compositions of the BC TME with patient survival [44], but to our knowledge, the current study is the first-of-its-kind to analyze the cell-type-specific transcriptomics profiles in the BC TME to quantify their respective contributions to treatment response prediction.

We applied DECODEM to investigate the predictive power of the expression of various cell types in the BC TME to patient response to NAC. Our findings highlight the prominence of immune cells (myeloid, plasmablasts and B-cells) in chemotherapy response, consistent with Kester et al. [44], elucidating the complementary roles of diverse cell types that allows for the incorporation of top predictive cell types to build accurate clinical response predictors, and showcase DECODEMi’s ability to identify key CCIs impacting response across HER2-BC. Furthermore, we illustrated DECODEM’s generalizability to SC transcriptomics capturing treatment-invariant properties of the TME, thereby enabling accurate classification of responders for chemotherapy and ICB. Moreover, we demonstrated DECODEMi’s ability to identify the top CCIs relevant to chemotherapy response and their generalizability to SC.

Immune cells have a clear association with prognosis in BC, both without and post chemotherapy in the early-stage setting [107][30][31]. The International TILs Working Group was established for the express purpose of standardizing incorporation of tumor infiltrating lymphocyte pathological review for clinical use and established an online prognosis calculator for patients with TNBC (https://www.tilsinbreastcancer.org/prognosis-tool/). Plasma cells and naïve B-cells have been associated with better pCR rates in ER+ tumors, whereas regulatory T-cells have been associated with better pCR rates in TNBC [108]. In our analysis, immune cells such as plasmablasts and myeloid cells exhibit involvement in signal transduction, particularly through VEGF and NOTCH signaling pathways, which are associated with tumor-associated angiogenesis, enrichment of BC stem cells and epithelial-to-mesenchymal transition, thus leading to poor prognosis and therapy resistance [61][62][65][67]. Stromal cells such as endothelial cells display enrichments in development- and immune-related pathways including L1CAM interactions and antigen processing and presentation, with L1CAM’s oncogenic role and defects in antigen processing leading to poor outcomes and therapy resistance [68][69][70][71][72]. Overall, our results paired with the existing clinical evidence reinforce that the presence of an active immune infiltration within the TME prior to treatment, as indicated by the prominent contribution of immune cells, leads to favorable chemotherapy response, as suggested by Sammut et al. [11].

DECODEM and DECODEMi complement the existing application of gene expression in the early-stage BC setting. Currently, gene expression-based tests such as Oncotype DX and MammaPrint are used for prediction and prognostication in ER+ patients, reporting if a subset should receive chemotherapy due to worse predicted outcomes [109]. These tests use quantitative RT-PCR or cDNA microarray to assess a known subset of genes and largely apply only to a subset of early-stage ER+ patients. DECODEM and DECODEMi provide complimentary context to these tools, expanding applicable subtypes for chemotherapy response prediction and the greater biological context.

Although we presented a comprehensive framework to assess the TME in a quantitative manner, certain limitations should be acknowledged and addressed in future studies. First, our predictors rely on data-driven algorithms involving deconvolution and ML, necessitating a large sample size to attain statistical robustness and avoid overfitting. Accordingly, the available cohorts are limited in size, limiting our ability to perform robust subtype-specific analyses. Second, any limitations inherent to CODEFACS (e.g., discrepancies in RNA-seq processing and reliance on appropriate molecular signatures [45]) will affect DECODEM in downstream analysis. Most of the cell types analyzed have well-defined gene signatures that allow for confident deconvolution and downstream analysis. However, it is known to be more challenging to distinguish between normal and cancer epithelial cells, which often employs copy number variation (CNV) inference methods that are highly dependent on the correlation between scRNA-seq and CNV along with the use of appropriate normal reference cells [110][111][112]. The same goes for DECODEMi and its reliance on LIRICS using an appropriate ligand – receptor database [45]. Third, our deconvolution analysis could not be performed separately for CD4 and CD8 T-cell subsets. Alternatively, we provided an enrichment analysis assessing the predictive powers of CD4 and CD8 T-cells that corroborated our findings from using the overall T-cells.

Our work focused on examining the association of the BC TME, specifically in HER2-BC, to clinical response to NAC. However, both DECODEM and DECODEMi can be applied to investigate diverse treatment regimens (as shown with ICB) in different cancer indications. Although both TransNEO and ARTemis + PBCP cohorts include different chemotherapy combinations, limited sample size prevented us from exploring further. Therefore, an intriguing direction could be to investigate the distinct role of the TME in different chemotherapy families with different mechanisms of actions (MOAs) e.g., taxanes, antimetabolites, platinum agents and so on. With DECODEMi, one could further pinpoint the family-specific significant CCIs to better elucidate the underlying MOA and uncover novel CCIs with promising therapeutic implications. Additionally, expanding the investigation beyond the scope of cancer and examining the impact of the microenvironment in various indications such as aging-related disorders and autoimmune diseases could provide further valuable insights.

In summary, our study introduces two data-driven computational frameworks that enable systematic investigation of the roles of the microenvironment in various processes, including treatment response. These frameworks can utilize bulk transcriptomics from healthy and diseased conditions when single-cell information is not easily accessible, providing a valuable approach to study the TME in a comprehensive manner. Their application in breast cancer uncovers the considerable predictive power of diverse cell types within the TME and highlights key stromal – immune CCIs that are strongly predictive of patient response. Taken together, we show that ensemble models integrating the expression of different cell types perform best and outperform models built on the original tumor bulk expression.

## METHODS

### Overview of the cohorts

We analyzed three bulk transcriptomics cohorts, encompassing a total of 255 breast cancer patients who received neoadjuvant chemotherapy (Figure 1A-B). The first cohort, recently published by Sammut et al., was the TransNEO cohort which included 94 HER2-breast tumors from patients undergoing various NAC regimens (Figure 1B) [11]. Notably, Sammut et al. presented the complete TransNEO cohort, covering 168 patients with early and locally advanced BC from all major subtypes, and provided DNA-seq, RNA-seq, digital pathology and treatment information for various subsets of patients [11]. The second cohort was the ARTemis + PBCP cohort, externally validated by Sammut et al., combining the control arm of the ARTemis trial (n = 38) received the combination therapy of FEC-T [47] and the HER2-cases from the Personalised Breast Cancer Programme (PBCP; n = 17) undergoing various NAC regimens [11] (Figure 1B). The third cohort, BrighTNess, included 106 patients from arm B of the BrighTNess trial, a randomized, double-blind, placebo-controlled three-arm clinical study for veliparib in stage II/III TNBC [48]. Arm B included only patients who received the NAC combination of paclitaxel, carboplatin, and a placebo for veliparib (Figure 1B). In all three cohorts, patient response was assessed at surgery using the residual cancer burden (RCB) index where patients achieving pathological complete response (pCR; RCB = 0) were taken as responders (R) and patients with residual diseases (RCB-I, II, III) as non-responders (NR) with different R:NR ratios (Figure 1A).

Furthermore, we analyzed two recent single-cell cohorts encompassing 21 patients with triple negative breast cancer, treated either with chemotherapy alone (n = 6) or chemotherapy and immune checkpoint blockade therapy (n = 15; Figure 1C). The first SC cohort was presented by Zhang et al. that consisted of 15 patients with pre-treatment tumor biopsy expression of four immune cell types (B-cells, innate lymphoid cells, myeloid and T-cells) and either treated with paclitaxel alone (n = 6) or combined with the anti-PD-L1 agent, atezolizumab (n = 9) [49]. Notably, this study originally comprised of 22 patients with advanced TNBC, where only half of the patients received combination therapy [49], and included 78 tumor biopsies and blood samples collected at three time-points including baseline. The patient response at surgery were assessed using the Response Evaluation Criteria in Solid Tumors (RECIST) criteria [49], where we categorized patients achieving complete or partial response as responders (R) and those with stable or progressive diseases as non-responders (NR). We divide Zhang et al. into two sub-cohorts for analysis based on treatment i.e., chemotherapy (n = 6) and chemotherapy + ICB (n = 9; Figure 1C). The second SC cohort was presented by Bassez et al. that consisted of 6 TNBC patients with pre-treatment tumor biopsy expression of six cell types (B-cells, CAFs, endothelial, malignant, myeloid and T-cells) and treated with taxane – anthracycline based neoadjuvant chemotherapy before receiving the anti-PD1 agent, pembrolizumab [50] (Figure 1C). This cohort was part of the window-of-opportunity study, BioKey, involving early diagnosed BC (including all major subtypes) that included one cohort of 29 treatment-naïve patients and another cohort of 11 patients treated with neoadjuvant chemotherapy (and anti-HER2 therapy for HER2+ patients) for 20-24 weeks, both followed by pembrolizumab before surgery. This study collected tumor biopsies at both pre- and on-treatment and performed scRNA-seq and scTCR-seq on the same SC suspension using 10x sequencing technologies [50]. As patient response were not available, we considered the post-treatment T-cell clonotype expansion assessed from scTCR-seq data as a surrogate variable while only considering the TNBC patients from the second cohort (n = 6), since they were treated with NAC and ICB therapy and included both clonotype expanders and non-expanders (E:NE = 3:3; unlike the 4 ER+ patients who displayed NE only) [50].

### Deconvolution with CODEFACS

The first step of DECODEM involves employing CODEFACS, a cellular deconvolution tool recently developed in our laboratory [45] (Figure 1D). CODEFACS characterizes the TME by reconstructing the cell-type-specific expression across the whole transcriptome in each sample from input bulk expression by utilizing the corresponding cell abundance or a cell-type-specific signature matrix as another input. The output includes the cell-type-specific expression profiles, cell abundance and a confidence score matrix quantifying the confidence level (between 0 and 1) of the inferred expression values for each gene in each cell type across all samples.

As a sanity check. we first tested the ability of CODEFACS to accurately deconvolve bulk expression profile into different cell-type-specific expression profiles for breast cancer samples. We leveraged a SC cohort recently presented by Wu et al. [46] that contained 26 primary breast tumors (11 ER+, 5 HER2+ and 10 TNBC) encompassing 130,246 single cells across nine major cell types and generated five benchmark cohorts (BC1 – BC5) following the benchmark generation procedure in Wang et al. [45]. BC1 contains pseudobulk expression for deconvolution and the corresponding ground truth data (cell abundance and cell-type-specific expression) for 22 out of 26 original patients (keeping 4 patients aside for signature generation with CIBERSORTx [105]); BC2 and BC3 contain 100 ‘pseudopatients’ each with different mixtures of cells; and BC4 and BC5 denote two noisy versions of BC1. Using CODEFACS, we inferred nine different cell types from each cohort and evaluated deconvolution accuracy by measuring the Kendall correlation (τ) between the observed and inferred values following Wang et al. [45], yielding high accuracy for both the cell-type abundance (Supplementary Figure 1A) and cell-type-specific expression (Supplementary Figure 1B). We also quantified thousands of accurately inferred genes (τ ≥ 0.3, P ≤ 0.05) on average across cell types (Supplementary Figure 1C).

We then used this scRNA-seq data (with original cell type annotation) from Wu et al. to derive a breast cancer-specific cell type signature matrix encompassing 1,400 genes representing the nine cell types using an in-house signature derivation tool based on CIBERSORTx [105]. We employed this signature matrix to deconvolve each bulk expression profile into nine cell-type-specific expression profiles across the unique protein-coding genes available per cohort (TransNEO = 17,680, ARTemis + PBCP = 17,940 and BrighTNess = 15,203). The aforementioned nine cell types were: B-cells, cancer-associated fibroblasts (CAFs), cancer epithelial (malignant), endothelial, myeloid, normal epithelial, plasmablasts, perivascular-like cells (PVL) and T-cells.

### Curation of a database of cell-type-specific ligand – receptor interactions in BC TME from single-cell

To infer the cell-cell interaction activity using LIRICS, we first constructed a database of cell-type-specific ligand – receptor interactions (LRIs) to characterize all cell-to-cell communications that are likely to occur within the TME of BC patients. To this end, we obtained a list of 2,422 potential L-R pairs annotated by Ramilowski et al. [51] and evaluated the potential of each pair occurring between the nine cell types of interest using the scRNA-seq data from Wu et al. again [46]. To discern the activity of the curated L-R pairs between cell types, we pooled together all 130,246 cells from Wu et al. and applied an empirical P-value based CCI inference approach using SOCIAL from Sahni et al. [75] (with a modified assumption of true LRI activity < null LRI activity in step 3), where an LRI with P ≤ 0.05 was categorized as ‘likely’ to occur in BC TME and ‘unlikely’ otherwise, following the standard practices in the field. This procedure enabled us to construct a database of 49,397 cell-type-specific L-R pairs (i.e., CCI quadruplets including ligand cell type, ligand gene, receptor cell type and receptor gene) that are likely to occur in the BC TME. This database composed the initial feature space for LIRICS to yield different numbers of CCIs across the three bulk cohorts (i.e., TransNEO = 46,755, ARTemis + PBCP = 43,887 and BrighTNess = 38,285).

### Extraction of cell-cell interactions with LIRICS

We extended DECODEM to DECODEMi by incorporating multi-cell-type information through CCIs (Figure 1D). To this end, DECODEMi applies LIRICS [45] on the output of CODEFACS to infer the corresponding CCI activity. LIRICS uses a two-step approach on deconvolved expression to infer the cell-type-specific ligand – receptor (L-R) pairs that are likely to be ‘active’ in each sample: First, it queries a database of 49,397 plausible L-R interactions between each pair of cell types in the BC TME, constructed using SC data (see above); Second, it then infers an L-R interaction as active in a pair of cell types in a given sample (indicated by ‘1’) if the expressions of both ligand and receptor genes exceed their median cell-type-specific expression values or inactive otherwise (indicated by ‘0’) [45]. We used this binary CCI activity matrix from LIRICS to assess the predictive potential of cell-to-cell communications.

### Machine learning pipeline with deconvolution output

The second step of both DECODEM and DECODEMi involves building ML predictors of clinical response using either cell-type-specific expression profiles or CCIs, respectively. For DECODEM, we built nine cell-type-specific predictors to classify R vs. NR following treatment using a four-stage ML pipeline (Figure 1D): (a) unsupervised prefiltering to keep only the genes that are deconvolved with a confidence score ≥ 0.99, (b) low-variance filtering to keep genes with variance ≥ 0.1, (c) supervised feature selection to select the top ‘m’ genes maximizing performance, where 2 ≤ m ≤ 25 with genes ranked by their ANOVA F-scores for association with response, followed by standardization to scale each gene to zero mean and unit variance, and (d) classification with an unweighted ensemble classifier comprised of regularized logistic regression, random forest, support vector machine and XGBoost (extreme gradient boosting) to predict the likelihood scores of response. The upper limit on the feature size is chosen to avoid model overfitting (by ensuring a lower feature size than sample size), reduce noise and facilitate shorter runtimes. Of note, we tested different feature sizes (m = 25, 50, 100) that yielded similar performance during model development and hence opted for the smallest size to ensure greater robustness. Furthermore, each score vector was rescaled between 0 and 1 across samples to improve out-of-distribution generalization. We trained the pipeline using three-fold cross-validation (CV) for hyperparameter tuning to maximize the area under the receiver operating characteristics (ROC) curve (AUC) for each classifier separately. To avoid overfitting, we repeated the training procedure five times with five different CV splits and took the mean of the five prediction vectors as our final prediction. Moreover, we investigated model performance by replacing the CV with a leave-one-out CV for hyperparameter tuning (maximizing accuracy) and obtained similar performance (Supplementary Figure 2), thereby opting to use the three-fold CV for main analyses. Once trained, we applied the test data to the trained models to yield model predictions and evaluate their performance using one-tailed Wilcoxon rank-sum test (P-value ≤ 0.05), ROC curve and precision-recall (PR) curve analysis. We further evaluate performance by computing the clinically relevant measure of diagnostic odds ratio (DOR), where we binarized the prediction scores using an optimal threshold learned from optimizing the training PR curve.

To accommodate for the sparse nature of SC transcriptomics (and CCI profile) for external validation, we slightly modified the pipeline to enable modeling for all available genes post filtering in step (c) i.e., 2 ≤ m ≤ ‘all’. Furthermore, to align SC expression with CODEFACS output (which provides the mean cell-type-specific expression profile per patient [45]), we harmonized the cell-type-specific SC expression by averaging across all corresponding cells for each patient, while only taking the top 3,500 highly variable genes to reduce noise. Additionally, for CCI-based analysis with DECODEMi, we skipped step (a) and further updated the low-variance cut-off in step (b) to 0.08 and the classifier in step (d) to random forest that is better suited to the binary nature of input CCI matrix and includes a built-in measure of feature importance, mean decrease in Gini impurity. For signed feature importance, the CCI directionalities were computed by using a two-tailed Fisher’s test (followed by false discovery rate (FDR) adjustment for multiple hypothesis correction i.e., FDR-adjusted P-value ≤ 0.05), where an odds ratio (OR) > 1 denotes an enrichment in responders and vice versa. Moreover, for filtering CCIs for the prominent cell types, we only include the interactions involving the confident L-R genes (CODEFACS confidence level ≥ 0.99) in the corresponding cell type.

For baseline comparison in DECODEM, we employed the same ML pipeline while skipping the prefiltering in step (a) for training with bulk transcriptomics (and further skipping feature selection in step (c) for training with cell abundance). For DECODEMi, we calculated the overall ‘pseudobulk’ expression (akin to the bulk mixture) as the mean expression across all cells covering all available cell types for each patient in a cohort post low-variance filtering (as described above). Additionally, since our framework dynamically adapts to the feature size available in both training and validation, we provide a validation script that generates the cell-type-specific and CCI-based models for a new validation cohort, essentially providing a ‘cohort-specific locked-down model’.

### CD4^+^ / CD8^+^ T-cells enrichment analysis

To analyze the contribution of the individual T-cell subtypes to chemotherapy response, we performed a gene set variation analysis (GSVA) to summarize the pathway activities per sample for each T-cell subtype. We employed two SC expression signatures derived from Wu et al. [46], collected via the Tumor Immune Single Cell Hub 2 (TISCH2) [106] and performed GSVA on the deconvolved T-cells expression using the GSVA R package. These signatures consisted of 207 and 424 differentially expressed genes (DEGs) for CD4 and CD8 T-cells, respectively, with log-fold-change ≥ 0.5 and an overlap of 151 DEGs. Subsequently, we evaluated the association of GSVA enrichment scores (rescaled between 0 and 1) with clinical response across the three bulk cohorts using a one-tailed Wilcoxon rank-sum test (FDR-adjusted P-value ≤ 0.05; Supplementary Figure 4A,C), ROC and precision-recall curve analyses (Supplementary Figure 4B,D).

### Cell-type-specific enrichment analysis

We conducted gene set enrichment analysis (GSEA) to investigate the association of each cell type with different biological processes. For each cell type, we first ranked the 17,680 protein-coding genes available in TransNEO by their association with positive outcome (i.e., pCR), measured by the P-values from a one-tailed Wilcoxon rank-sum test, adjusted by CODEFACS confidence scores. We performed a separate GSEA analysis using the Reactome pathways [53] for each prominent cell type (except CAFs), yielding six different sets of enriched pathways (FDR-adjusted P-value ≤ 0.2) across six prominent cell types. From these sets, we reported the top 10 relevant pathways for each cell type (Figure 3G).

### Single-cell validation of top CCIs from bulk

We validated the most predictive CCIs extracted by DECODEMi in bulk using a validation cohort sourced from the pre-treatment SC expression of six TNBC patients from Zhang et al., who were treated with NAC [49]. We devised a three-step validation procedure: First, for consistency, we retrained DECODEMi with TransNEO including only the interactions among cell types available in both bulk and SC (B-cells, myeloid and T-cells). This enabled us to extract the top predictive CCIs (m = 170) involving a total of 134 unique L-R pairs. Second, to enhance the statistical power and robustness, we pooled the 28,209 cells encompassing the three cell types across all six patients and randomly downsampled 30% cells 100 times within each response group for each cell type. This procedure generated a profile of 200 pseudopatients (R:NR = 100:100) with empirical P-values denoting the significance levels of the 804 possible CCIs involving the 134 L-R pairs in three cell types (= 134 × 2 × 3), yielding the ‘SC-TNBC cohort’. To identify the activation status of these 170 CCIs of interest within this cohort, we applied the SOCIAL-based CCI inference approach [75] described above to denote a CCI status as ‘active’ if P ≤ 0.05 and ‘inactive’ otherwise. Third, for each pseudopatient, we predicted chemotherapy response as the sum of active CCIs, signed by their directionalities (i.e., + or - when OR > 1 or < 1, FDR-adjusted P-value ≤ 0.05; and 0 otherwise), followed by a rescaling between 0 and 1 across all pesudopatients. We then used these prediction scores to validate the performance of these top CCIs using a one-tailed Wilcoxon rank-sum test (FDR-adjusted P-value ≤ 0.05) and ROC curve analysis (Figure 4G-H).

### Patient survival stratification in TCGA-BRCA

We explored whether the DECODEM scores can be used to stratify the survival of the 484 early-stage HER2-breast cancer patients (encompassing stages I – IIIA following the training cohort, TransNEO, that included early-stage BC patients only) in The Cancer Genome Atlas (TCGA-BRCA) from their cell-type-specific expression. Focusing on the expression of the prominent cell type of malignant cells (top performer across subtypes; Supplementary Figure 3), DECODEM effectively stratified the patients within each BC subtype into High-score and Low-score groups (implying low-and high-risk, respectively) using a fixed threshold of 0.5, as displayed by their Kaplan-Meier curves for overall survival (Log-rank test P-value ≤ 0.05, Supplementary Figure 8A-B) and progression-free interval (Supplementary Figure 9A-B). The remaining prominent cell types also displayed similar survival trends (Supplementary Figure 8C-N, Supplementary Figure 9C-N).

## DATA AVAILABILITY

For TransNEO cohort, RNA-seq counts and clinical data were acquired from the publication by Sammut et al. (https://doi.org/10.1038/s41586-021-04278-5). RNA-seq counts were converted to TPM by using Ensembl annotations (GRCh37.87, http://ftp.ensembl.org/pub/grch37/release-87/gtf/homo_sapiens/). For ARTemis + PBCP cohort, clinical data were collected from Sammut et al. (https://doi.org/10.1038/s41586-021-04278-5) and RNA-seq TPM values were obtained from the authors. For BrighTNess cohort, RNA-seq FPKM values (converted to TPM) and clinical data were acquired from the gene expression omnibus (GEO; accession no: GSE164458; https://www.ncbi.nlm.nih.gov/geo/query/acc.cgi?acc=GSE164458). For Zhang et al., scRNA-seq counts and clinical data were obtained from the publication itself (https://doi.org/10.1016/j.ccell.2021.09.010). For Bassez et al., scRNA-seq counts and clinical data were obtained from Lambrechts Lab website (https://lambrechtslab.sites.vib.be/en/single-cell). For Wu et al., scRNA-seq counts and cell annotations were collected from GEO (accession no: GSE176078; https://www.ncbi.nlm.nih.gov/geo/query/acc.cgi?acc=GSE176078) and the list of differentially expressed genes across cell types were collected from TISCH2 (http://tisch.comp-genomics.org/gallery/). The ligand-receptor list from Ramilowski et al. were collected via GitHub (https://github.com/LewisLabUCSD/Ligand-Receptor-Pairs). For TCGA-BRCA, RNA-seq counts (converted to TPM), survival data and clinical data were collected from the UCSC Xena browser (https://xenabrowser.net). All deconvolved expression and cell-cell interaction data generated in this study will be made available upon publication.

## CODE AVAILABILITY

All relevant codes for reproducing the figures in this study are publicly available upon publication at https://github.com/ruppinlab/DECODEM/. The codes for CODEFACS and LIRICS are available from the original publication at https://zenodo.org/record/5790343/. The codes for molecular signature generation are available at https://github.com/ruppinlab/scSigR/.

## ACKNOWLEDGEMENTS

This research is supported in part by the Intramural Research Program of the National Institutes of Health, National Cancer Institute, Center for Cancer Research. This work has utilized the computational resources of the NIH HPC Biowulf cluster (http://hpc.nih.gov). The results presented are in part based on the data generated by The Cancer Genome Atlas Research network (https://www.cancer.gov/tcga/). We would additionally like to acknowledge Thomas Cantore for helpful discussion regarding cellular deconvolution and survival analysis, Sanna Madan and Danh-Tai Hoang for guidance pertaining cell type identification and data visualization, E. Michael Gertz for data collection and helpful discussion regarding survival analysis, and the members of the Cancer Data Science Laboratory for their constructive feedback. Panel D in Figure 1 has been created in BioRender (Dhruba, S. R. (2025) https://BioRender.com/z38y774). ChatGPT v4o (https://chatgpt.com/) was used strictly to refine certain existing paragraphs for better overall presentation.

## AUTHOR CONTRIBUTIONS

S.R.D, K.W. and E.R. conceived and designed the study. S.R.D., S.S. and K.W. performed the model development and validation analyses. S.R.D., K.W., S.S., P.S.R., B.W., Y.S., E.S., S.Si., S-J.S., C.C. and E.R. acquired, analyzed or interpreted the data. S.R.D., S.S., K.W., D.W., S.Si. and B.W. worked on data visualization. E.R. and K.W. supervised the study. All authors critically revised the manuscript for important intelligent content.

---

**Supplementary Figure 1:**
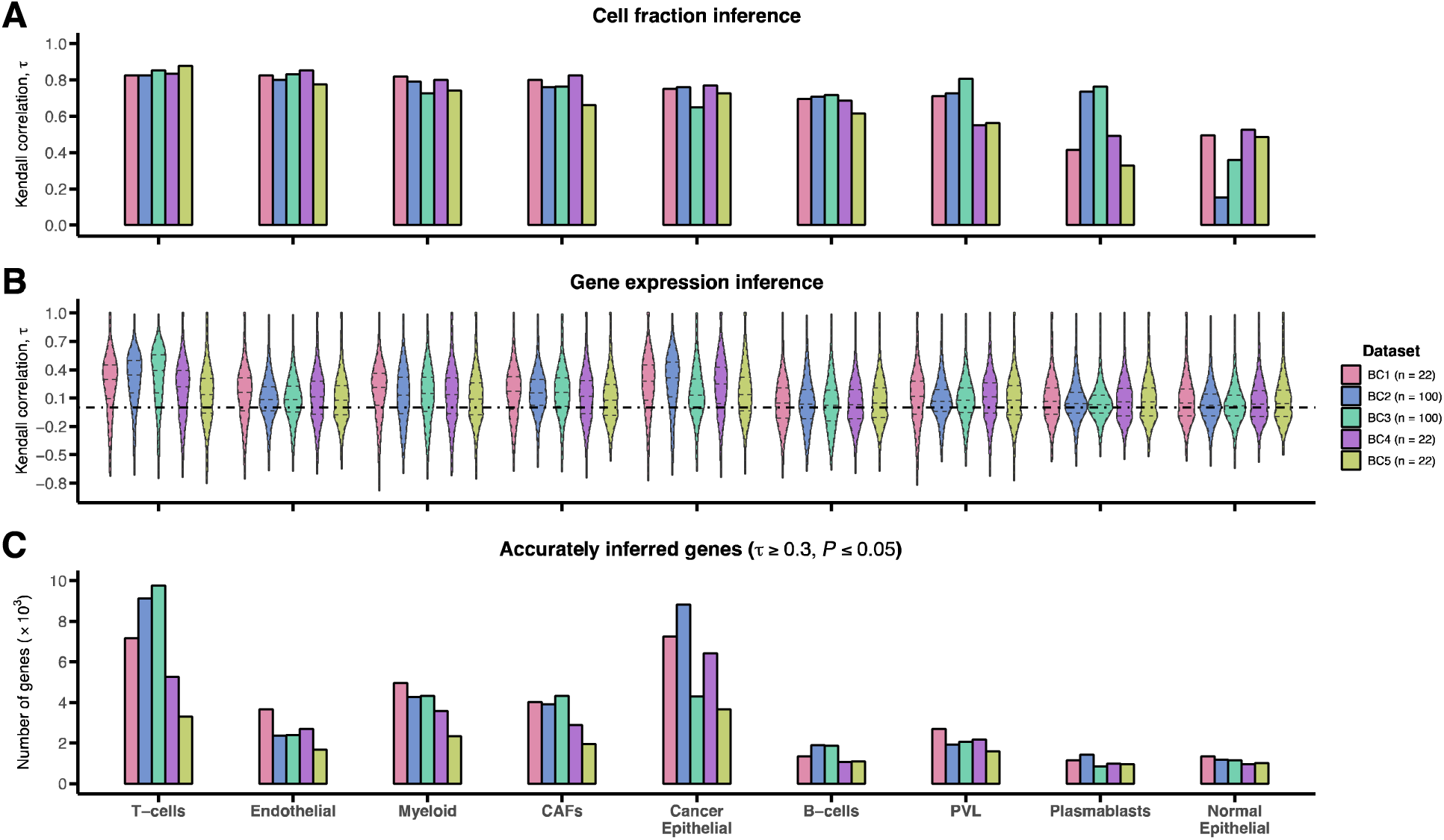
Benchmarking CODEFACS on five breast cancer cohorts generated from single-cell (SC) data. **A.** Correlation of actual and CODEFACS-inferred cell fraction values for nine cell types across five benchmark cohorts (BC1-5), generated via mixing SC expression from Wu et al. (see METHODS). Cell types are ranked by their mean correlation across the five cohorts in a descending order. **B.** Correlation of actual and CODEFACS-inferred cell-type-specific gene expression values for nine cell types across five benchmark cohorts. Cell types are ranked by their mean correlation for cell abundance accuracy in A. **C.** Number of accurately inferred genes for nine cell types across five benchmark cohorts. Cell types are ranked by their mean correlation for cell abundance accuracy in A.

**Supplementary Figure 2:**
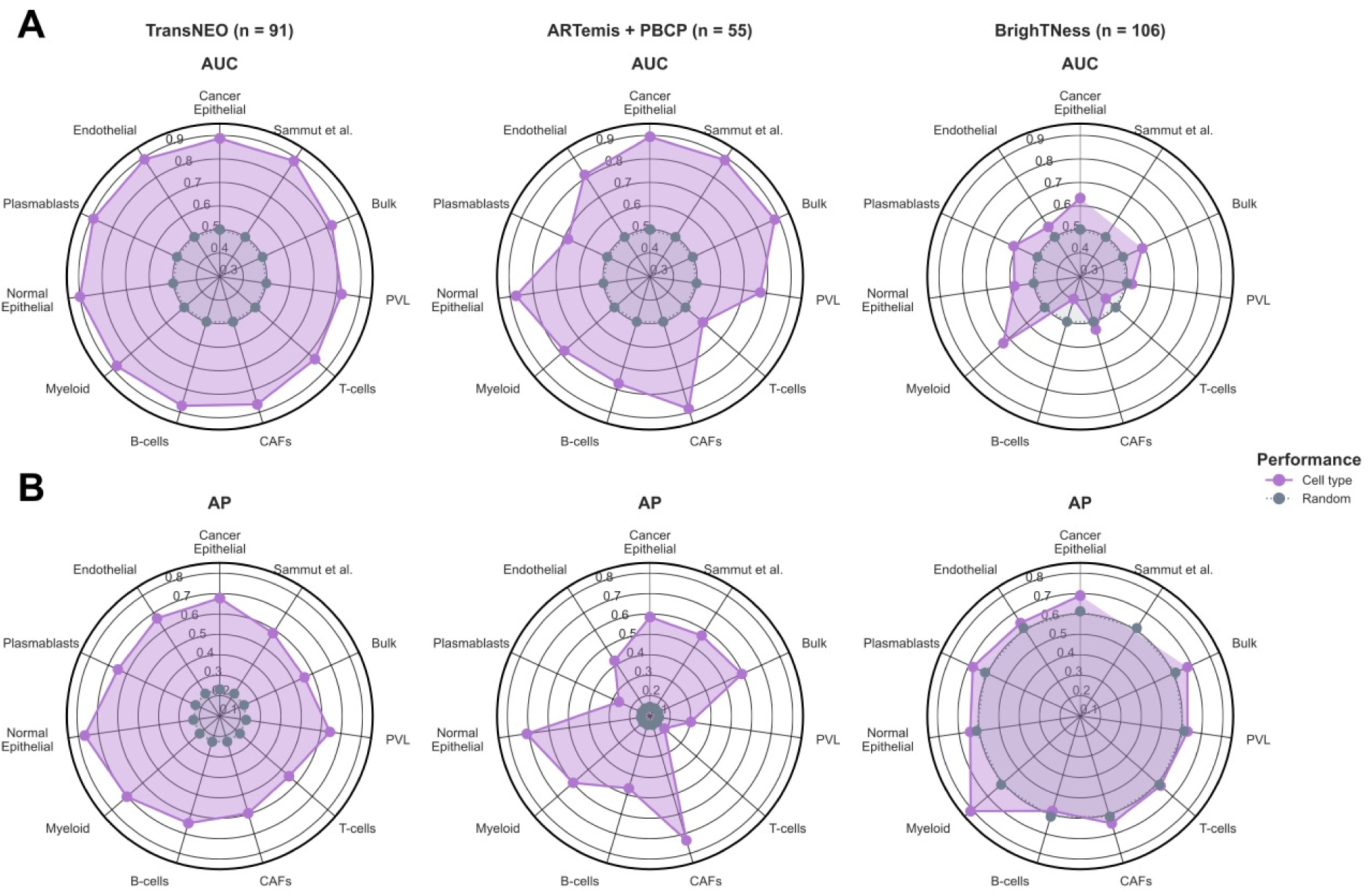
Hyperparameter tuning with leave-one-out cross-validation (CV) leads to similar cell-type-specific performance to using three-fold cross-validation. **A-B.** Comparison of model performance for nine cell-type-specific, bulk and Sammut et al. (for first two cohorts) predictors across TransNEO, ARTemis + PBCP and BrighTNess, where model hyperparameters were optimized using leave-one-out CV. AUC and AP stand for the area under the receiver operating characteristics curve and average precision, respectively. ‘Random’ denotes a random predictor (AUC = 0.5, AP = fraction of responders). Cell types are ranked by their original AUC values for TransNEO (obtained using three-fold CV for hyperparameter tuning; see Figure 2A) in the counterclockwise direction.

**Supplementary Figure 3:**
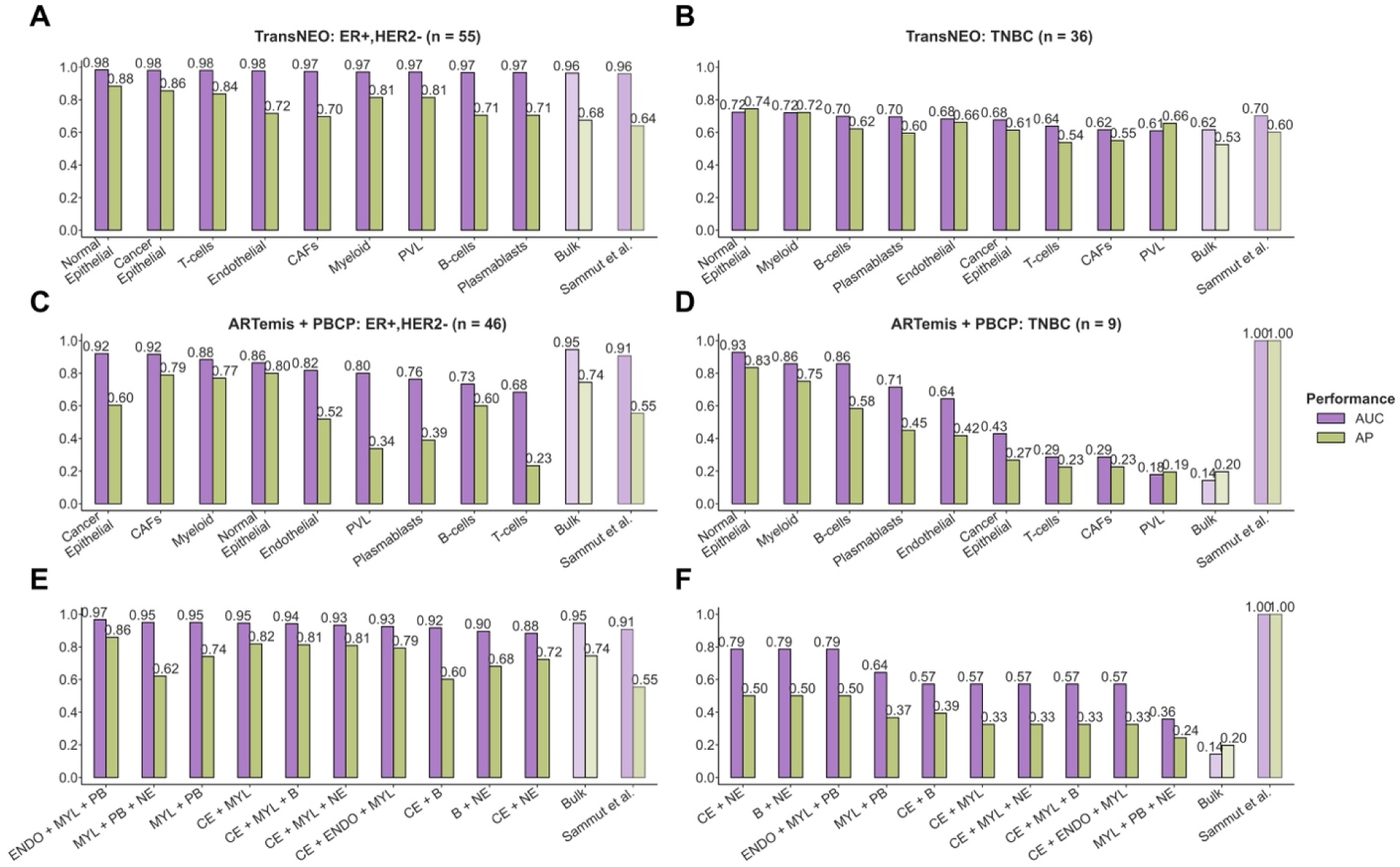
The prominent cell types and ensembles mediating chemotherapy response in breast cancer tumor microenvironment in a subtype-specific manner. **A-B.** Comparison of model performance across TransNEO for the ER+,HER2- (A) and TNBC (B) subtypes for nine cell-type-specific, bulk and Sammut et al. predictors. AUC and AP stand for the area under the receiver operating characteristics curve and average precision (equivalent to the area under the precision-recall curve), respectively. Cell types are ranked by their AUC values in a descending order. **C-D.** Comparison of model performance across ARTemis + PBCP for the ER+,HER2- (C) and TNBC (**D**) subtypes for nine cell-type-specific, bulk and Sammut et al. predictors. Cell types are ranked by their AUC values in a descending order. **E-F.** Comparison of model performance across with ARTemis + PBCP for the ER+,HER2- (E) and TNBC (F) subtypes for the five most prominent two-cell-ensembles, five most prominent three-cell-ensembles, bulk and Sammut et al. predictors. Ensembles are ranked by their AUC values in a descending order.

**Supplementary Figure 4:**
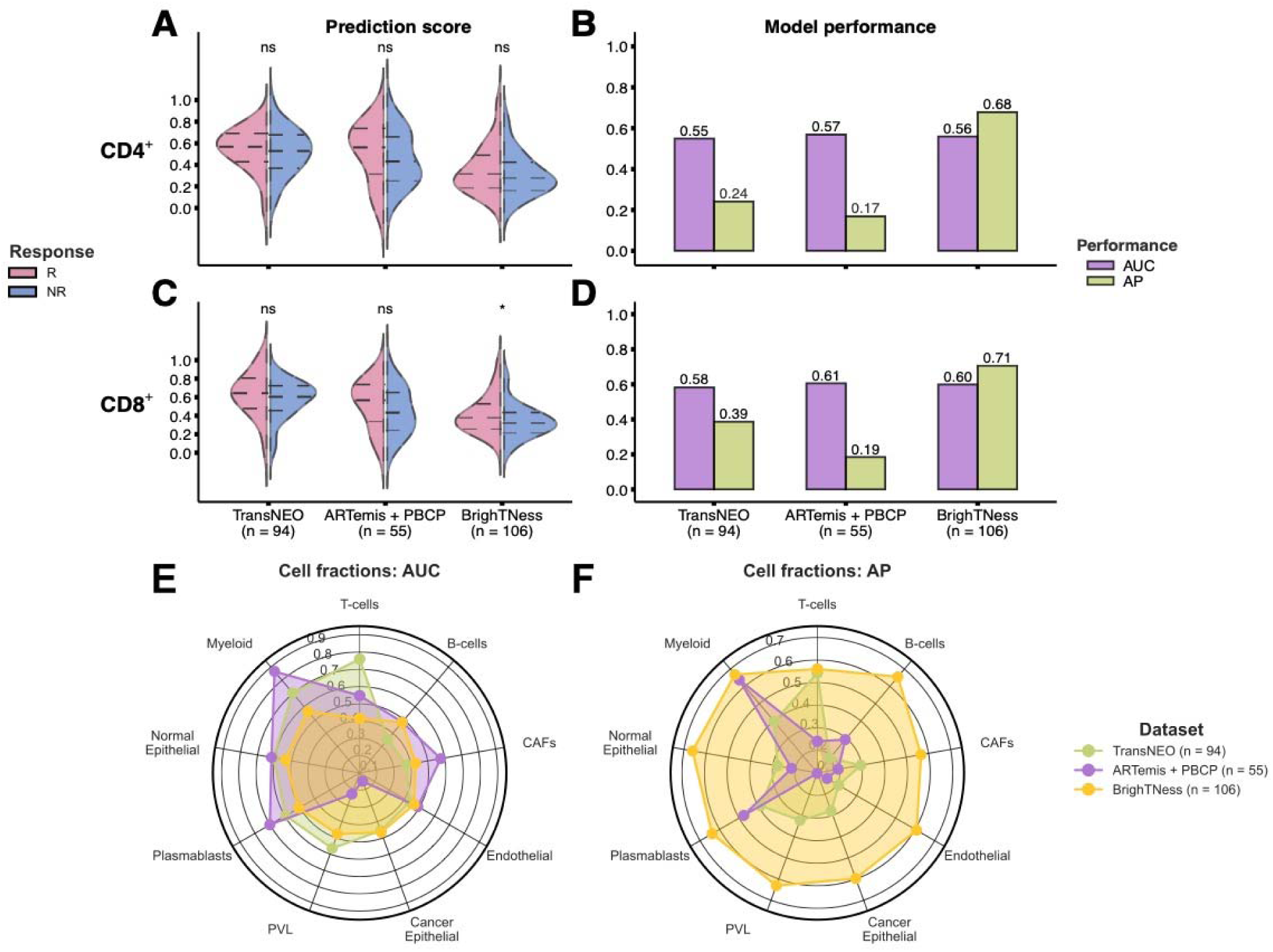
The CD4 / CD8 T-cells enrichment and cell abundance – clinical response association analyses reaffirm the poor chemotherapy response stratification by T-cells at the current resolution. **A-B.** GSVA enrichment scores (A) and predictive performance (B) of CD4 T-cells for chemotherapy response stratification across TransNEO, ARTemis + PBCP and BrighTNess. R and NR stand for responders and non-responders, respectively. The differences between the prediction scores were computed by using a one-tailed Wilcoxon rank-sum test (* and ‘ns’ denote P ≤ 0.05 and P > 0.05, respectively). AUC and AP stand for the area under the receiver operating characteristics curve and average precision (equivalent to the area under precision-recall curve), respectively. **C-D.** GSVA enrichment scores (C) and predictive performance (D) of CD8 T-cells for chemotherapy response stratification across the three cohorts. **E-F.** Predictive performance of abundances of the nine cell types across the three cohorts. Cell types are ranked by their AUC values for TransNEO in a descending order in the counterclockwise direction.

**Supplementary Figure 5:**
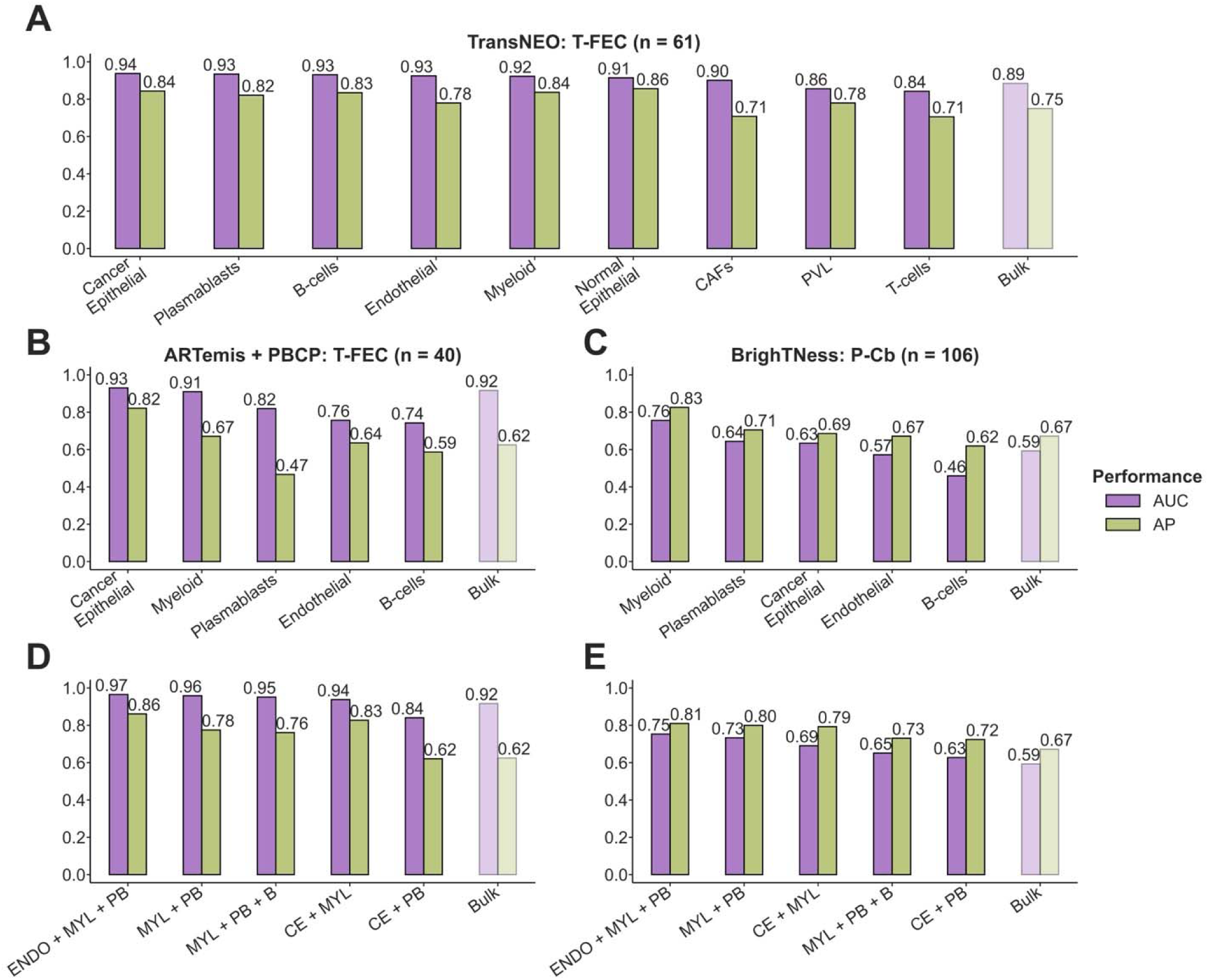
The prominent immune cell types and ensembles mediating response to the most prevalent immunogenic chemotherapy regimen in each breast cancer cohort. **A.** Comparison of model performance for nine cell-type-specific and bulk predictors across TransNEO for the T-FEC regimen. AUC and AP stand for the area under the receiver operating characteristics curve and average precision (equivalent to the area under the precision-recall curve), respectively. C, E, F and T stand for cyclophosphamide, epirubicin, 5-flurouracil and docetaxel (taxotere), respectively. Cell types are ranked by their AUC values in a descending order. **B-C.** Comparison of model performance for the top five cell-type-specific and bulk predictor across ARTemis + PBCP for the T-FEC regiment (B) and BrighTNess for the paclitaxel – carboplatin (P-Cb) regimen (C). Cell types are ranked by their AUC values in a descending order. D-E. Comparison of model performance for the top five multi-cell-type ensemble and bulk predictors across ARTemis + PBCP for the T-FEC regiment (D) and BrighTNess for the P-Cb regimen (E). Ensembles are ranked by their AUC values in a descending order.

**Supplementary Figure 6:**
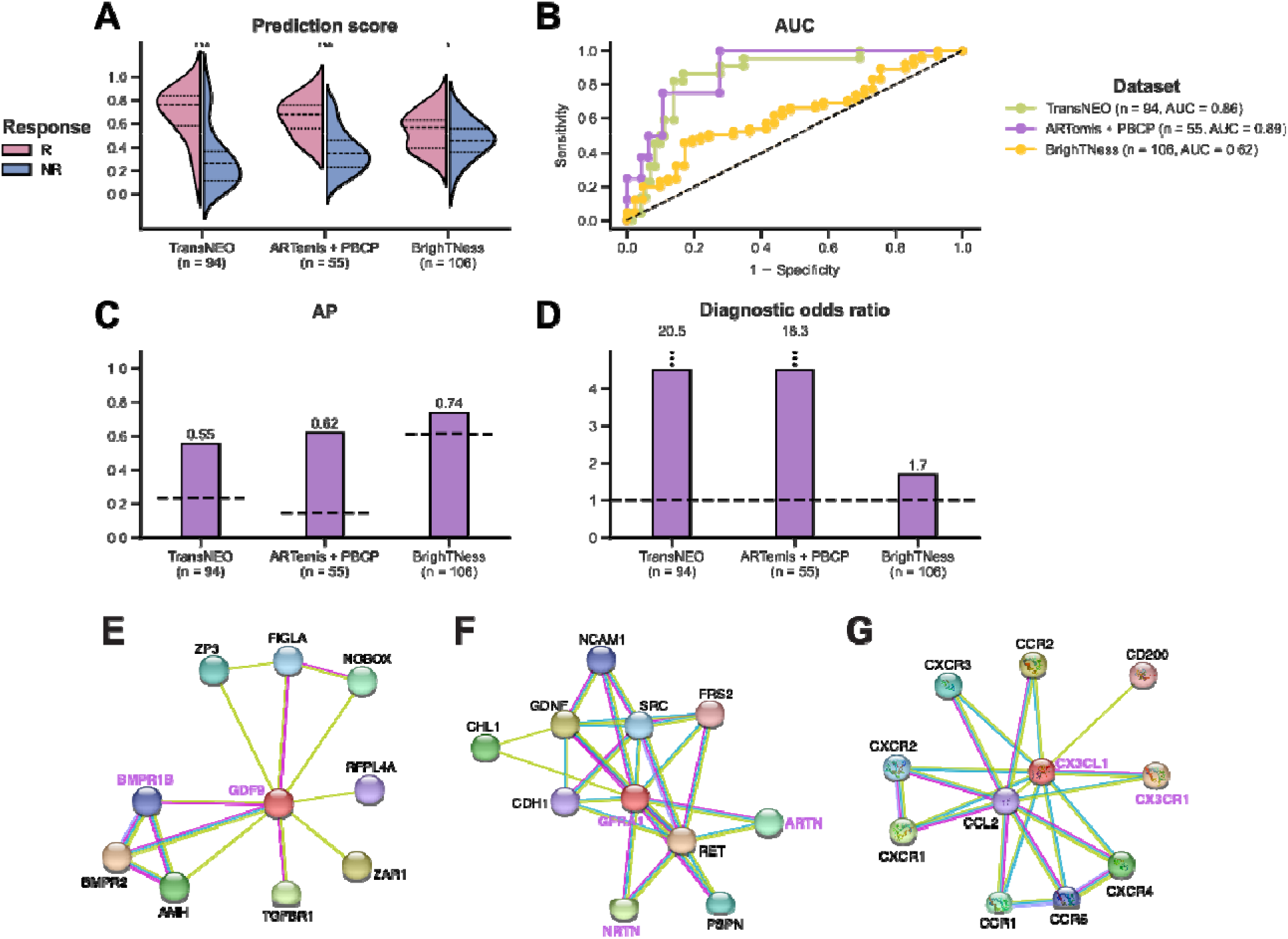
Cell-cell interactions (CCIs) encompassing all cell types in mediating chemotherapy response in breast cancer tumor microenvironment. **A-D.** Comparison of prediction scores (A) and model performance (B-D) for DECODEMi CCI-based predictor using all available CCIs across TransNEO, ARTemis + PBCP and BrighTNess. R and NR stand for responders and non-responders, respectively. The differences between the prediction scores were computed by using a one-tailed Wilcoxon rank-sum test (*** and * denote P ≤ 0.001 and P ≤ 0.05, respectively). AUC and AP stand for the area under the receiver operating characteristics curve and average precision, respectively. The dotted lines in C-D represent the AP and diagnostic odds ratio (DOR) values for a random predictor (AP = fraction of responders, DOR = 1.0). **E-G.** Protein-protein interaction networks obtained from STRING [76] displaying the known interaction networks for three prominent CCIs i.e., GDF9 – BMPR1B (E), NRTN – GFRA1 and ARTN – GFRA1 (F) and CX3CL1 – CX3CR1 (G).

**Supplementary Figure 7:**
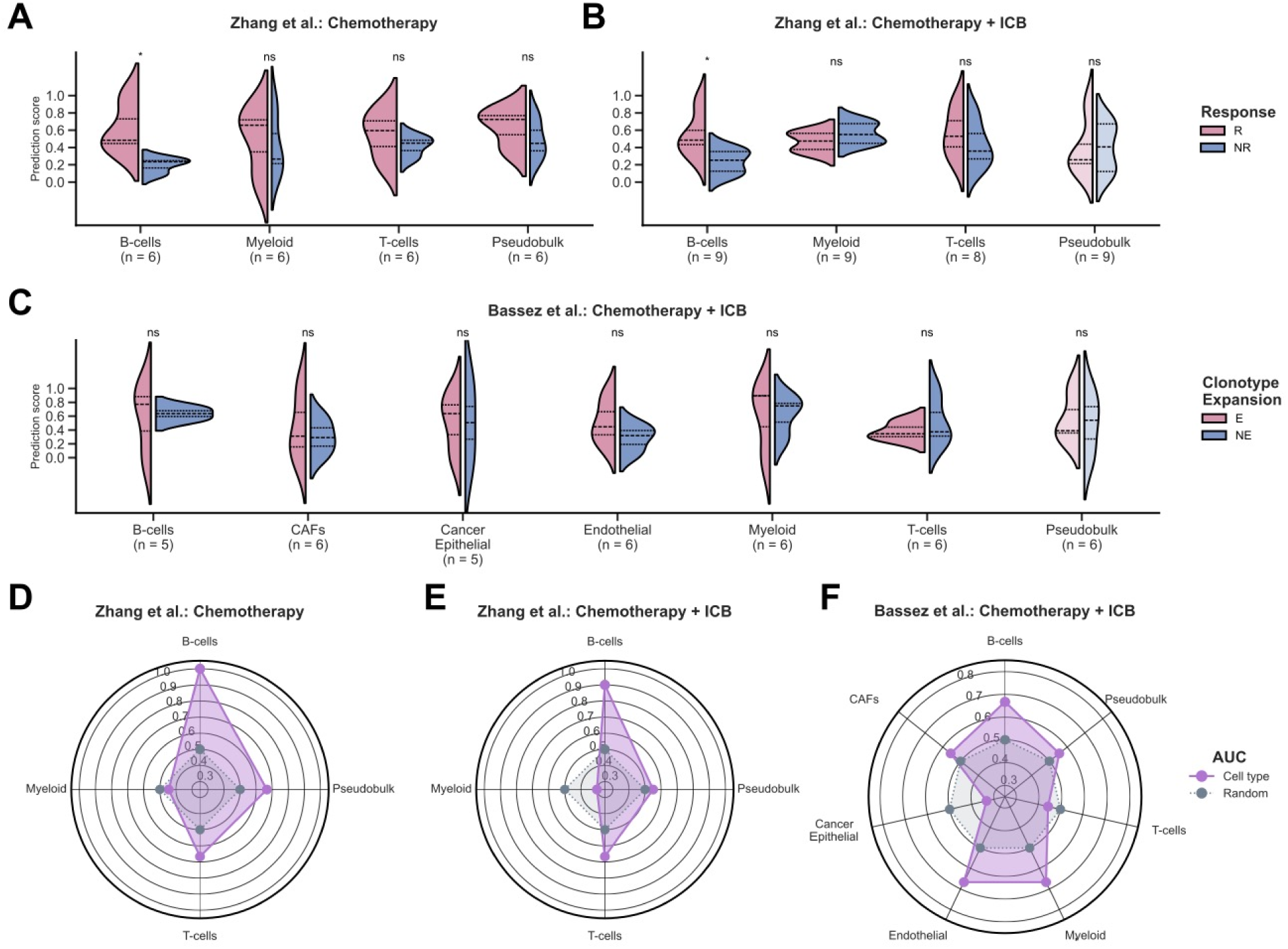
DECODEM generalizes to single-cell (SC) transcriptomics for patient response prediction. **A-B.** Comparison of prediction scores for the cell-type-specific and pseudobulk (similar to bulk in previous scenarios) predictors across SC expression for predicting response to neoadjuvant chemotherapy (NAC) alone (A) or in combination with immune checkpoint blockade (ICB) therapy (B) from Zhang et al., or predicting T-cell clonotype expansion following treatment with NAC and ICB therapy (C) from Bassez et al. R, NR, E and NE stand for responders, non-responders, expanders and non-expanders, respectively. The differences between the prediction scores were computed by using a one-tailed Wilcoxon rank-sum test (* and ‘ns’ denote P ≤ 0.05 and P > 0.05, respectively). **D-F.** Comparison of model performance for the cell-type-specific and pseudobulk predictors for predicting response to NAC alone (D) or in combination with ICB therapy (E) from Zhang et al., and for predicting T-cell clonotype expansion following treatment with NAC and ICB therapy from Bassez et al. (F). AUC stand for the area under the receiver operating characteristics curve. ‘Random’ denotes a random predictor (AUC = 0.5).

**Supplementary Figure 8:**
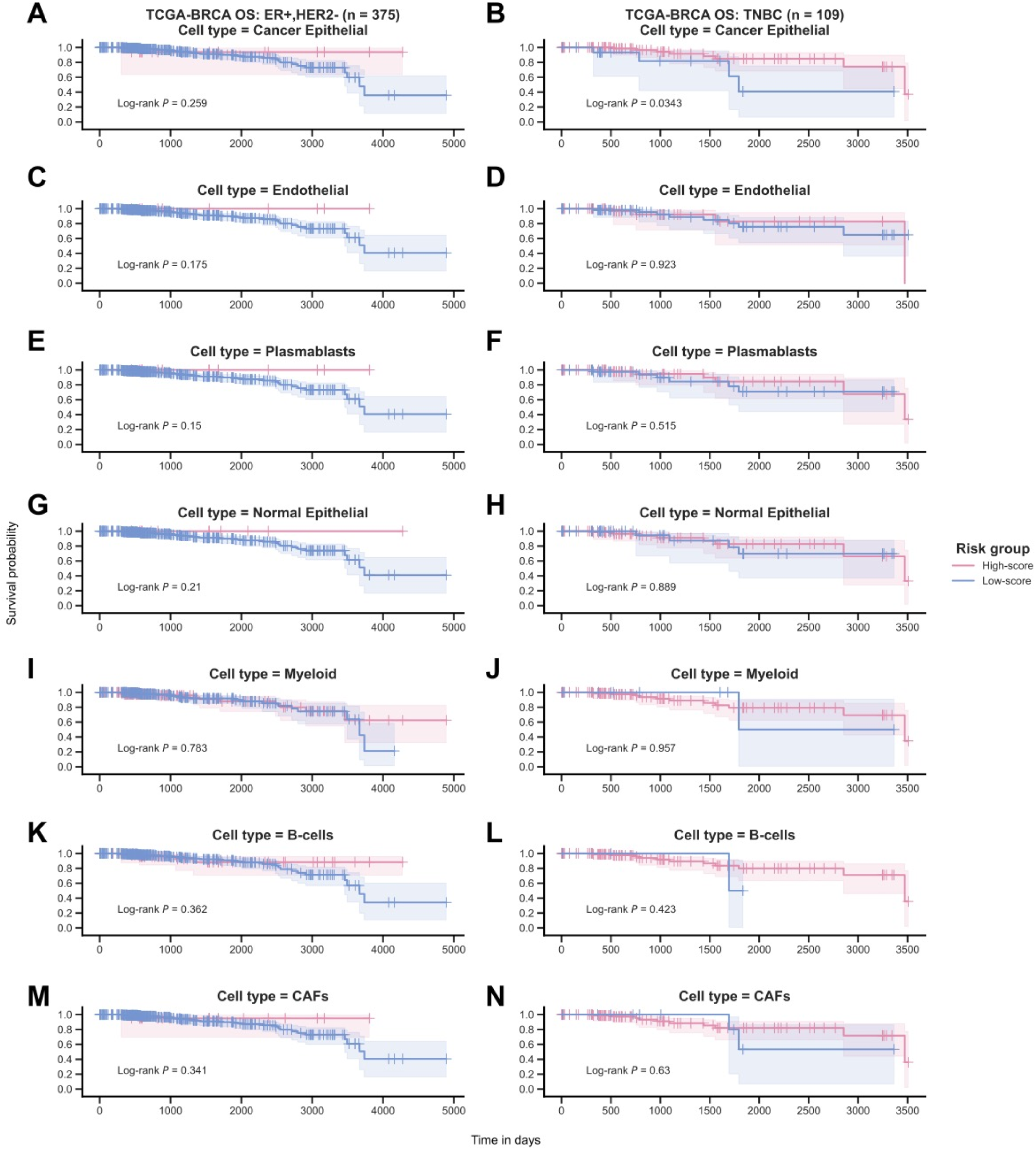
The generalizability of DECODEM to the stratification of TCGA-BRCA overall survival (OS). **A-N.** Kaplan-Meier curves depicting OS of early stage TCGA-BRCA patients with ER+,HER2-BC (left) and TNBC (right), stratified by DECODEM scores for seven prominent cell types. Patients were stratified into ‘High-score’ and ‘Low-score’ groups using a threshold of 0.5 on their DECODEM scores for these seven cell types. The differences between the curves were computed by using a Log-rank test.

**Supplementary Figure 9:**
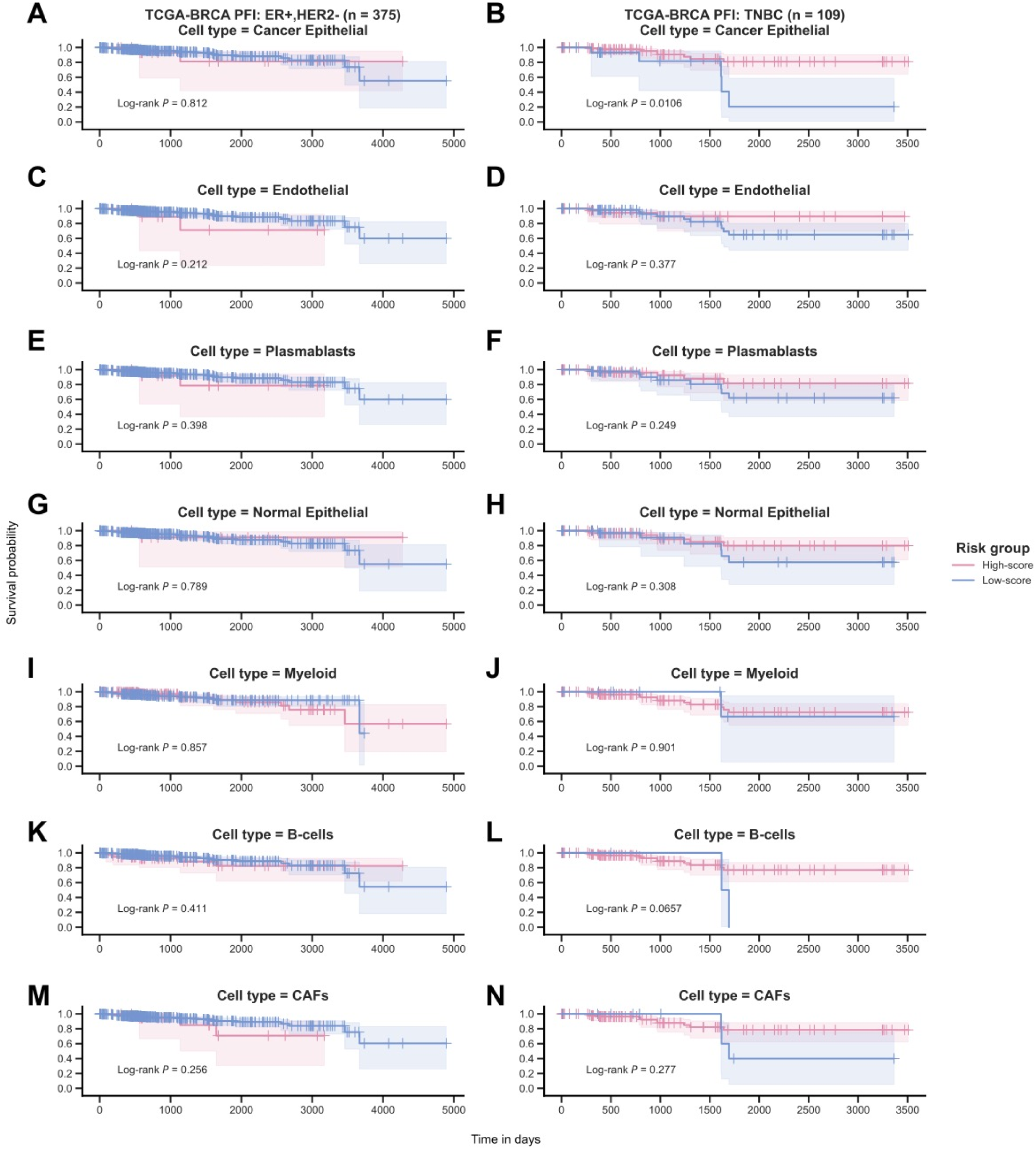
The generalizability of DECODEM to the stratification TCGA-BRCA progression-free interval (PFI; synonymous to progression-free survival). **A-N.** Kaplan-Meier curves depicting PFI of early stage TCGA-BRCA patients with ER+,HER2-BC (left) and TNBC (right), stratified by DECODEM scores for seven prominent cell types. Patients were stratified into ‘High-score’ and ‘Low-score’ groups using a threshold of 0.5 on their DECODEM scores for these seven cell types. The differences between the curves were computed by using a Log-rank test.

## REFERENCES

[1] Tsimberidou, A. M., Fountzilas, E., Nikanjam, M., & Kurzrock, R. (2020). Review of precision cancer medicine: Evolution of the treatment paradigm. Cancer treatment reviews, 86, 102019.

[2] Huang K, Xiao C, Glass LM, Critchlow CM,. Machine learning applications for therapeutic tasks with genomics data. Patterns 2021; 2(10):100328.

[3] Fisher, B., Brown, A., Mamounas, E., Wieand, S., Robidoux, A., Margolese, R. G., … & Dimitrov, N. V. (1997). Effect of preoperative chemotherapy on local-regional disease in women with operable breast cancer: findings from National Surgical Adjuvant Breast and Bowel Project B-18. Journal of clinical oncology, 15(7), 2483–2493.

[4] Bear, H. D., Anderson, S., Brown, A., Smith, R., Mamounas, E. P., Fisher, B., … & Wolmark, N. (2003). The effect on tumor response of adding sequential preoperative docetaxel to preoperative doxorubicin and cyclophosphamide: preliminary results from National Surgical Adjuvant Breast and Bowel Project Protocol B-27. Journal of Clinical Oncology, 21(22), 4165–4174.

[5] Golshan, M., Loibl, S., Wong, S. M., Huober, J. B., O’Shaughnessy, J., Rugo, H. S., … & Untch, M. (2020). Breast conservation after neoadjuvant chemotherapy for triple-negative breast cancer: surgical results from the BrighTNess randomized clinical trial. JAMA surgery, 155(3), e195410–e195410.

[6] Urueña, C., Lasso, P., Bernal-Estevez, D. et al. The breast cancer immune microenvironment is modified by neoadjuvant chemotherapy. Sci Rep 12, 7981 (2022). 10.1038/s41598-022-12108-5

[7] Ali, H. R., Dariush, A., Provenzano, E., Bardwell, H., Abraham, J. E., Iddawela, M., … & Caldas, C. (2016). Computational pathology of pre-treatment biopsies identifies lymphocyte density as a predictor of response to neoadjuvant chemotherapy in breast cancer. Breast Cancer Research, 18(1), 1–11.

[8] Ali, H. R., Dariush, A., Thomas, J., Provenzano, E., Dunn, J., Hiller, L., … & Caldas, C. (2017). Lymphocyte density determined by computational pathology validated as a predictor of response to neoadjuvant chemotherapy in breast cancer: secondary analysis of the ARTemis trial. Annals of Oncology, 28(8), 1832–1835.

[9] Dieci, M. V., Prat, A., Tagliafico, E., Paré, L., Ficarra, G., Bisagni, G., … & Guarneri, V. (2016). Integrated evaluation of PAM50 subtypes and immune modulation of pCR in HER2-positive breast cancer patients treated with chemotherapy and HER2-targeted agents in the CherLOB trial. Annals of Oncology, 27(10), 1867–1873.

[10] Griguolo, G., Serna, G., Pascual, T. et al. Immune microenvironment characterisation and dynamics during anti-HER2-based neoadjuvant treatment in HER2-positive breast cancer. npj Precis. Onc. 5, 23 (2021). HYPERLINK ”10.1038/s41698-021-00163-6. 10.1038/s41698-021-00163-64040

[11] Sammut, SJ., Crispin-Ortuzar, M., Chin, Sf. et al. Multi-omic machine learning predictor of breast cancer therapy response. Nature 601, 623–629 (2022). 10.1038/s41586-021-04278-5

[12] Klemm F, Joyce JA. Microenvironmental regulation of therapeutic response in cancer. Trends Cell Biol. 2015 Apr;25(4):198–213. doi: 10.1016/j.tcb.2014.11.006. Epub 2014 Dec 22. PMID: 25540894; PMCID: PMC5424264.

[13] Son B, Lee S, Youn H, Kim E, Kim W, Youn B. The role of tumor microenvironment in therapeutic resistance. Oncotarget. 2017 Jan 17;8(3):3933–3945. doi: 10.18632/oncotarget.13907. PMID: 27965469; PMCID: PMC5354804.

[14] Wu, P., Gao, W., Su, M., Nice, E. C., Zhang, W., Lin, J., & Xie, N. (2021). Adaptive mechanisms of tumor therapy resistance driven by tumor microenvironment. Frontiers in cell and developmental biology, 9, 641469.

[15] Mehraj, U., Dar, A.H., Wani, N.A. et al. Tumor microenvironment promotes breast cancer chemoresistance. Cancer Chemother Pharmacol 87, 147–158 (2021). 10.1007/s00280-020-04222-w

[16] Ruffell, B., & Coussens, L. M. (2015). Macrophages and therapeutic resistance in cancer. Cancer cell, 27(4), 462–472.

[17] Acharyya, S., Oskarsson, T., Vanharanta, S., Malladi, S., Kim, J., Morris, P. G., … & Massagué, J. (2012). A CXCL1 paracrine network links cancer chemoresistance and metastasis. Cell, 150(1), 165–178.

[18] Li, M., Quintana, A., Alberts, E., Hung, M. S., Boulat, V., Ripoll, M. M., & Grigoriadis, A. (2023). B Cells in Breast Cancer Pathology. Cancers, 15(5), 1517.

[19] Sakaguchi, A., Horimoto, Y., Onagi, H. et al. Plasma cell infiltration and treatment effect in breast cancer patients treated with neoadjuvant chemotherapy. Breast Cancer Res 23, 99 (2021). 10.1186/s13058-021-01477-w

[20] Yeong, J., Lim, J. C. T., Lee, B., Li, H., Chia, N., Ong, C. C. H., … & Iqbal, J. (2018). High densities of tumor-associated plasma cells predict improved prognosis in triple negative breast cancer. Frontiers in immunology, 9, 1209.

[21] Bhinder, B., Gilvary, C., Madhukar, N. S., & Elemento, O. (2021). Artificial intelligence in cancer research and precision medicine. Cancer Discovery, 11(4), 900–915.

[22] Singla, N., & Singla, S. (2020). Harnessing big data with machine learning in precision oncology. Kidney cancer journal: official journal of the Kidney Cancer Association, 18(3), 83.

[23] Senft, D., Leiserson, M. D., Ruppin, E., & Ze’ev, A. R. (2017). Precision oncology: the road ahead. Trends in molecular medicine, 23(10), 874–898.

[24] Tsimberidou, A. M., Fountzilas, E., Bleris, L., & Kurzrock, R. (2020, September). Transcriptomics and solid tumors: The next frontier in precision cancer medicine. In Seminars in cancer biology. Academic Press.

[25] Siravegna, G., Marsoni, S., Siena, S., & Bardelli, A. (2017). Integrating liquid biopsies into the management of cancer. Nature reviews Clinical oncology, 14(9), 531–548.

[26] Heitzer, E., Haque, I. S., Roberts, C. E., & Speicher, M. R. (2019). Current and future perspectives of liquid biopsies in genomics-driven oncology. Nature Reviews Genetics, 20(2), 71–88.

[27] Sawabata, N. (2020). Circulating tumor cells: From the laboratory to the cancer clinic. Cancers, 12(10), 3065.

[28] Beaubier, N., Bontrager, M., Huether, R., Igartua, C., Lau, D., Tell, R., … & White, K. P. (2019). Integrated genomic profiling expands clinical options for patients with cancer. Nature biotechnology, 37(11), 1351–1360.

[29] Hayashi, H., Takiguchi, Y., Minami, H., Akiyoshi, K., Segawa, Y., Ueda, H., … & Nakagawa, K. (2020). Site-specific and targeted therapy based on molecular profiling by next-generation sequencing for cancer of unknown primary site: a nonrandomized phase 2 clinical trial. JAMA oncology, 6(12), 1931–1938.

[30] Vaske, O. M., Bjork, I., Salama, S. R., Beale, H., Shah, A. T., Sanders, L., … & Haussler, D. (2019). Comparative tumor RNA sequencing analysis for difficult-to-treat pediatric and young adult patients with cancer. JAMA network open, 2(10), e1913968–e1913968.

[31] Lee, J. S., Nair, N. U., Dinstag, G., Chapman, L., Chung, Y., Wang, K., … & Ruppin, E. (2021). Synthetic lethality-mediated precision oncology via the tumor transcriptome. Cell, 184(9), 2487–2502.

32. Sinha, S., Dhruba, S. R., Wu, W., Kerr, D. L., Stroganov, O. V., Grishagin, I., … & Ruppin, E. (2022). Predicting patient treatment response and resistance via single-cell transcriptomics of their tumors. bioRxiv.

[33] Dinstag, G., Shulman, E. D., Elis, E., Ben-Zvi, D. S., Tirosh, O., Maimon, E., … & Aharonov, R. (2022). Clinically oriented prediction of patient response to targeted and immunotherapies from the tumor transcriptome. bioRxiv.

[34] Fu, Y., Jung, A. W., Torne, R. V., Gonzalez, S., Vöhringer, H., Shmatko, A., … & Gerstung, M. (2020). Pan-cancer computational histopathology reveals mutations, tumor composition and prognosis. Nature Cancer, 1(8), 800–810.

[35] Noorbakhsh, J., Farahmand, S., Namburi, S., Caruana, D., Rimm, D., Soltanieh-ha, M., … & Chuang, J. H. (2020). Deep learning-based cross-classifications reveal conserved spatial behaviors within tumor histological images. Nature communications, 11(1), 1–14.

[36] Yu, K. H., Wang, F., Berry, G. J., Re, C., Altman, R. B., Snyder, M., & Kohane, I. S. (2020). Classifying non-small cell lung cancer types and transcriptomic subtypes using convolutional neural networks. Journal of the American Medical Informatics Association, 27(5), 757–769.

[37] Couture, H. D., Williams, L. A., Geradts, J., Nyante, S. J., Butler, E. N., Marron, J. S., … & Niethammer, M. (2018). Image analysis with deep learning to predict breast cancer grade, ER status, histologic subtype, and intrinsic subtype. NPJ breast cancer, 4(1), 1–8.

[38] Qu, H., Zhou, M., Yan, Z., Wang, H., Rustgi, V. K., Zhang, S., … Metaxas, D. N. (2021). Genetic mutation and biological pathway prediction based on whole slide images in breast carcinoma using deep learning. NPJ precision oncology, 5(1), 1–11.

[39] Hoang, D. T., Dinstag, G., Hermida, L. C., Ben-Zvi, D. S., Elis, E., Caley, K., … & Ruppin, E. (2022). Synthetic lethality-based prediction of cancer treatment response from histopathology images. bioRxiv.

[40] Tanioka, M., Fan, C., Parker, J. S., Hoadley, K. A., Hu, Z., Li, Y., … & Perou, C. M. (2018). Integrated Analysis of RNA and DNA from the Phase III Trial CALGB 40601 Identifies Predictors of Response to Trastuzumab-Based Neoadjuvant Chemotherapy in HER2-Positive Breast CancerIntegrated Response Models in HER2-Positive Breast Cancer. Clinical Cancer Research, 24(21), 5292–5304.

[41] Wong, M., Mayoh, C., Lau, L., Khuong-Quang, D. A., Pinese, M., Kumar, A., … & Cowley, M.J. (2020). Whole genome, transcriptome and methylome profiling enhances actionable target discovery in high-risk pediatric cancer. Nature medicine, 26(11), 1742–1753.

[42] Rodon, J., Soria, J. C., Berger, R., Miller, W. H., Rubin, E., Kugel, A., … & Kurzrock, R. (2019). Genomic and transcriptomic profiling expands precision cancer medicine: the WINTHER trial. Nature medicine, 25(5), 751–758.

[43] Korde, L. A., Somerfield, M. R., Carey, L. A., Crews, J. R., Denduluri, N., Hwang, E. S., … & Hershman, D. L. (2021). Neoadjuvant chemotherapy, endocrine therapy, and targeted therapy for breast cancer: ASCO guideline. Journal of Clinical Oncology, 39(13), 1485–1505.

44. Kester, L., Seinstra, D., van Rossum, A. G., Vennin, C., Hoogstraat, M., van der Velden, D., … & van Rheenen, J. (2022). Differential Survival and Therapy Benefit of Patients with Breast Cancer Are Characterized by Distinct Epithelial and Immune Cell Microenvironments Tumor Cellular Composition Predicts Benefit to Therapies. Clinical Cancer Research, OF1-OF12.

[45] Wang, K., Patkar, S., Lee, J.S., Gertz, E.M., Robinson, W., Schischlik, F., Crawford, D.R., Schäffer, A.A. and Ruppin, E. (2022). Deconvolving Clinically Relevant Cellular Immune Cross-talk from Bulk Gene Expression Using CODEFACS and LIRICS Stratifies Patients with Melanoma to Anti–PD-1 Therapy. Cancer discovery, 12(4), 1088–1105.

[46] Wu, S. Z., Al-Eryani, G., Roden, D. L., Junankar, S., Harvey, K., Andersson, A., … & Swarbrick, A. (2021). A single-cell and spatially resolved atlas of human breast cancers. Nature genetics, 53(9), 1334–1347.

[47] Earl, H. M., Hiller, L., Dunn, J. A., Blenkinsop, C., Grybowicz, L., Vallier, A. L., … & Hayward, L. (2015). Efficacy of neoadjuvant bevacizumab added to docetaxel followed by fluorouracil, epirubicin, and cyclophosphamide, for women with HER2-negative early breast cancer (ARTemis): an open-label, randomised, phase 3 trial. The lancet oncology, 16(6), 656–666.

[48] Loibl, S., O’Shaughnessy, J., Untch, M., Sikov, W. M., Rugo, H. S., McKee, M. D., … & Geyer Jr, C. E. (2018). Addition of the PARP inhibitor veliparib plus carboplatin or carboplatin alone to standard neoadjuvant chemotherapy in triple-negative breast cancer (BrighTNess): a randomised, phase 3 trial. The Lancet Oncology, 19(4), 497–509.

[49] Zhang, Y., Chen, H., Mo, H., Hu, X., Gao, R., Zhao, Y., … & Liu, Z. (2021). Single-cell analyses reveal key immune cell subsets associated with response to PD-L1 blockade in triple-negative breast cancer. Cancer Cell, 39(12), 1578–1593.

50. Bassez, A., Vos, H., Van Dyck, L., Floris, G., Arijs, I., Desmedt, C., … & Lambrechts, D. (2021). A single-cell map of intratumoral changes during anti-PD1 treatment of patients with breast cancer. Nature medicine, 27(5), 820–832.

[51] Ramilowski, J., Goldberg, T., Harshbarger, J. et al. A draft network of ligand–receptor-mediated multicellular signalling in human. Nat Commun 6, 7866 (2015). 10.1038/ncomms8866

[52] Green, D. R., Ferguson, T., Zitvogel, L., & Kroemer, G. (2009). Immunogenic and tolerogenic cell death. Nature Reviews Immunology, 9(5), 353–363.

[53] Fabregat, A. et al. The Reactome Pathway Knowledgebase. Nucleic Acids Res. 46, D649– D655 (2018).

[54] Molinari, M. (2000). Cell cycle checkpoints and their inactivation in human cancer. Cell proliferation, 33(5), 261–274.

[55] Bower, J.J., Vance, L.D., Psioda, M. et al. Patterns of cell cycle checkpoint deregulation associated with intrinsic molecular subtypes of human breast cancer cells. npj Breast Cancer 3, 9 (2017). 10.1038/s41523-017-0009-7

[56] Zhou, K., Sun, Y., Dong, D. et al. EMP3 negatively modulates breast cancer cell DNA replication, DNA damage repair, and stem-like properties. Cell Death Dis 12, 844 (2021). 10.1038/s41419-021-04140-6

[57] Repo, H., Löyttyniemi, E., Kurki, S. et al. A prognostic model based on cell-cycle control predicts outcome of breast cancer patients. BMC Cancer 20, 558 (2020). 10.1186/s12885-020-07045-3

[58] Vihervuori, H., Korpinen, K., Autere, T. A., Repo, H., Talvinen, K., & Kronqvist, P. (2022). Varying outcomes of triple-negative breast cancer in different age groups–prognostic value of clinical features and proliferation. Breast Cancer Research and Treatment, 1-12.

[59] Thu, K. L., Soria-Bretones, I., Mak, T. W., & Cescon, D. W. (2018). Targeting the cell cycle in breast cancer: towards the next phase. Cell Cycle, 17(15), 1871–1885.

[60] Zhang, J., Chan, D. W., & Lin, S. Y. (2022). Exploiting DNA Replication Stress as a Therapeutic Strategy for Breast Cancer. Biomedicines, 10(11), 2775.

[61] Zhu, X., & Zhou, W. (2015). The emerging regulation of VEGFR-2 in triple-negative breast cancer. Frontiers in Endocrinology, 6, 159.

[62] Ceci, C., Atzori, M. G., Lacal, P. M., & Graziani, G. (2020). Role of VEGFs/VEGFR-1 signaling and its inhibition in modulating tumor invasion: Experimental evidence in different metastatic cancer models. International journal of molecular sciences, 21(4), 1388.

[63] Zhang, Q., Lu, S., Li, T., Yu, L., Zhang, Y., Zeng, H., … & Lin, Y. (2019). ACE2 inhibits breast cancer angiogenesis via suppressing the VEGFa/VEGFR2/ERK pathway. Journal of Experimental & Clinical Cancer Research, 38(1), 1–12.

[64] Edwards, A., & Brennan, K. (2021). Notch signalling in breast development and cancer. Frontiers in Cell and Developmental Biology, 9, 692173.

[65] Zhou, B., Lin, W., Long, Y. et al. Notch signaling pathway: architecture, disease, and therapeutics. Sig Transduct Target Ther 7, 95 (2022). 10.1038/s41392-022-00934-y.

[66] Miao, K., Lei, J.H., Valecha, M.V. et al. NOTCH1 activation compensates BRCA1 deficiency and promotes triple-negative breast cancer formation. Nat Commun 11, 3256 (2020). 10.1038/s41467-020-16936-9

[67] BeLow, M., & Osipo, C. (2020). Notch signaling in breast cancer: a role in drug resistance. Cells, 9(10), 2204.

[68] Li, L., Wang, X., Hu, K. et al. ZNF133 is a potent suppressor in breast carcinogenesis through dampening L1CAM, a driver for tumor progression. Oncogene (2023). 10.1038/s41388-023-02731-5

69. van der Maten, M., Reijnen, C., Pijnenborg, J. M., & Zegers, M. M. (2019). L1 cell adhesion molecule in cancer, a systematic review on domain-specific functions. International journal of molecular sciences, 20(17), 4180.

[70] Zhang, J., Yang, F., Ding, Y., Zhen, L., Han, X., Jiao, F., & Tang, J. (2015). Overexpression of L1 cell adhesion molecule correlates with aggressive tumor progression of patients with breast cancer and promotes motility of breast cancer cells. International journal of clinical and experimental pathology, 8(8), 9240.

[71] Bandola-Simon, J., & Roche, P. A. (2019). Dysfunction of antigen processing and presentation by dendritic cells in cancer. Molecular immunology, 113, 31–37.

[72] Dhatchinamoorthy, K., Colbert, J. D., & Rock, K. L. (2021). Cancer immune evasion through loss of MHC class I antigen presentation. Frontiers in immunology, 12, 636568.

[73] McDonnell, A. M., Lesterhuis, W. J., Khong, A., Nowak, A. K., Lake, R. A., Currie, A. J., & Robinson, B. W. (2015). Tumor-infiltrating dendritic cells exhibit defective cross-presentation of tumor antigens, but is reversed by chemotherapy. European journal of immunology, 45(1), 49–59.

[74] Galluzzi, L., Humeau, J., Buqué, A., Zitvogel, L., & Kroemer, G. (2020). Immunostimulation with chemotherapy in the era of immune checkpoint inhibitors. Nature reviews Clinical oncology, 17(12), 725–741.

[75] Sahni, S., Wang, B., Wu, D., Dhruba, S. R., Nagy, M., Patkar, S., … & Ruppin, E. (2024). A machine learning model reveals expansive downregulation of ligand-receptor interactions that enhance lymphocyte infiltration in melanoma with developed resistance to immune checkpoint blockade. Nature Communications, 15(1), 8867.

[76] Szklarczyk, D., Gable, A. L., Lyon, D., Junge, A., Wyder, S., Huerta-Cepas, J., … & Mering, C. V. (2019). STRING v11: protein–protein association networks with increased coverage, supporting functional discovery in genome-wide experimental datasets. Nucleic acids research, 47(D1), D607–D613.

[77] Li, J., Li, C., Liu, X. et al. GDF9 concentration in embryo culture medium is linked to human embryo quality and viability. J Assist Reprod Genet 39, 117–125 (2022). 10.1007/s10815-021-02368-x

[78] Stocker, W. A., Walton, K. L., Richani, D., Chan, K. L., Beilby, K. H., Finger, B. J., … & Harrison, C. A. (2020). A variant of human growth differentiation factor-9 that improves oocyte developmental competence. Journal of Biological Chemistry, 295(23), 7981–7991.

79. Gomez-Puerto, M. C., Iyengar, P. V., García de Vinuesa, A., Ten Dijke, P., & Sanchez-Duffhues, G. (2019). Bone morphogenetic protein receptor signal transduction in human disease. The journal of pathology, 247(1), 9–20.

[80] Feary, E. S., Juengel, J. L., Smith, P., French, M. C., O’Connell, A. R., Lawrence, S. B., … & McNatty, K. P. (2007). Patterns of expression of messenger RNAs encoding GDF9, BMP15, TGFBR1, BMPR1B, and BMPR2 during follicular development and characterization of ovarian follicular populations in ewes carrying the Woodlands FecX2W mutation. Biology of reproduction, 77(6), 990–998.

[81] Hanavadi, S., Martin, T. A., Watkins, G., Mansel, R. E., & Jiang, W. G. (2007). The role of growth differentiation factor-9 (GDF-9) and its analog, GDF-9b/BMP-15, in human breast cancer. Annals of surgical oncology, 14, 2159–2166.

[82] Harrath, A. H., Jalouli, M., Oueslati, M. H., Farah, M. A., Feriani, A., Aldahmash, W., … & Alwasel, S. (2021). The flavonoid, kaempferol-3-O-apiofuranosyl-7-O-rhamnopyranosyl, as a potential therapeutic agent for breast cancer with a promoting effect on ovarian function. Phytotherapy Research, 35(11), 6170–6180.

[83] Bach, D. H., Park, H. J., & Lee, S. K. (2018). The dual role of bone morphogenetic proteins in cancer. Molecular Therapy-Oncolytics, 8, 1–13.

[84] Bokobza, S. M., Ye, L., Kynaston, H. E., Mansel, R. E., & Jiang, W. G. (2009). Reduced expression of BMPR-IB correlates with poor prognosis and increased proliferation of breast cancer cells. Cancer genomics & proteomics, 6(2), 101–108.

[85] Zheng, Y., Jiang, X., Wang, M., Yang, S., Deng, Y., Li, Y., … & Chen, L. (2022). BMPR1B Polymorphisms (rs1434536 and rs1970801) are Associated With Breast Cancer Susceptibility in Northwest Chinese Han Females: A Case-Control Study. Clinical Breast Cancer, 22(5), e641–e646.

[86] Dai, K., Qin, F., Zhang, H., Liu, X., Guo, C., Zhang, M., … & Ma, Y. (2016). Low expression of BMPRIB indicates poor prognosis of breast cancer and is insensitive to taxane-anthracycline chemotherapy. Oncotarget, 7(4), 4770.

[87] Jing, S., Wen, D., Yu, Y., Holst, P. L., Luo, Y., Fang, M., … & Fox, G. M. (1996). GDNF–induced activation of the ret protein tyrosine kinase is mediated by GDNFR-α, a novel receptor for GDNF. Cell, 85(7), 1113–1124.

[88] Fielder, G. C., Yang, T. W. S., Razdan, M., Li, Y., Lu, J., Perry, J. K., … & Liu, D. X. (2018). The GDNF family: a role in cancer?. Neoplasia, 20(1), 99–117.

[89] Li, Q., Cao, Z., & Zhao, S. (2022). The Emerging Portrait of Glial Cell Line-derived Neurotrophic Factor Family Receptor Alpha (GFRα) in Cancers. International Journal of Medical Sciences, 19(4), 659.

[90] Pecar, G., Liu, S., Hooda, J. et al. RET signaling in breast cancer therapeutic resistance and metastasis. Breast Cancer Res 25, 26 (2023). 10.1186/s13058-023-01622-7

[91] Horibata, S., Rice, E. J., Mukai, C., Marks, B. A., Sams, K., Zheng, H., … & Danko, C. G. (2018). ER-positive breast cancer cells are poised for RET-mediated endocrine resistance. PLoS One, 13(4), e0194023.

[92] Bhakta, S., Crocker, L. M., Chen, Y., Hazen, M., Schutten, M. M., Li, D., … & Junutula, J. R. (2018). An Anti-GDNF Family Receptor Alpha 1 (GFRA1) Antibody–Drug Conjugate for the Treatment of Hormone Receptor–Positive Breast Cancer Targeting Hormone-Positive Breast Cancer with a GFRA1 ADC. Molecular cancer therapeutics, 17(3), 638–649.

[93] Kim, M., & Kim, D. J. (2018). GFRA1: a novel molecular target for the prevention of osteosarcoma chemoresistance. International journal of molecular sciences, 19(4), 1078.

[94] Liu, H., Yang, Z., Lu, W., Chen, Z., Chen, L., Han, S., … & Cai, Y. (2020). Chemokines and chemokine receptors: A new strategy for breast cancer therapy. Cancer medicine, 9(11), 3786–3799.

[95] Liang, Y., Yi, L., Liu, P., Jiang, L., Wang, H., Hu, A., … & Dong, J. (2018). CX3CL1 involves in breast cancer metastasizing to the spine via the Src/FAK signaling pathway. Journal of Cancer, 9(19), 3603.

[96] Tardáguila, M., Mira, E., García-Cabezas, M. A., Feijoo, A. M., Quintela-Fandino, M., Azcoitia, I., … & Manes, S. (2013). CX3CL1 Promotes Breast Cancer via Transactivation of the EGF PathwayCX3CL1 Promotes Breast Cancer. Cancer research, 73(14), 4461–4473.

[97] Jamieson-Gladney, W.L., Zhang, Y., Fong, A.M. et al. The chemokine receptor CX3CR1 is directly involved in the arrest of breast cancer cells to the skeleton. Breast Cancer Res 13, R91 (2011). 10.1186/bcr3016

[98] Rivas-Fuentes, S., Salgado-Aguayo, A., Arratia-Quijada, J., & Gorocica-Rosete, P. (2021). Regulation and biological functions of the CX3CL1-CX3CR1 axis and its relevance in solid cancer: A mini-review. Journal of Cancer, 12(2), 571.

[99] Yue, Y., Zhang, Q., & Sun, Z. (2022). CX3CR1 acts as a protective biomarker in the tumor microenvironment of colorectal Cancer. Frontiers in Immunology, 12, 5972.

[100] Ishida, Y., Kuninaka, Y., Yamamoto, Y., Nosaka, M., Kimura, A., Furukawa, F., … & Kondo, T. (2020). Pivotal involvement of the CX3CL1-CX3CR1 axis for the recruitment of M2 tumor-associated macrophages in skin carcinogenesis. Journal of Investigative Dermatology, 140(10), 1951–1961.

[101] Cortes, J., Haiderali, A., Huang, M., Pan, W., Schmid, P., Akers, K. G., … & O’Shaughnessy, J. (2023). Neoadjuvant immunotherapy and chemotherapy regimens for the treatment of high-risk, early-stage triple-negative breast cancer: a systematic review and network meta-analysis. BMC cancer, 23(1), 792.

[102] Schmid, P., Cortes, J., Pusztai, L., McArthur, H., Kümmel, S., Bergh, J., … & O’Shaughnessy, J. (2020). Pembrolizumab for early triple-negative breast cancer. New England Journal of Medicine, 382(9), 810–821.

[103] Kim, L., Coman, M., Pusztai, L., & Park, T. S. (2023). Neoadjuvant Immunotherapy in Early, Triple-Negative Breast Cancers: Catching Up with the Rest. Annals of Surgical Oncology, 30(11), 6441–6449.

[104] Michaud, M., Martins, I., Sukkurwala, A. Q., Adjemian, S., Ma, Y., Pellegatti, P., … & Kroemer, G. (2011). Autophagy-dependent anticancer immune responses induced by chemotherapeutic agents in mice. Science, 334(6062), 1573–1577.

[105] Newman, A. M., Steen, C. B., Liu, C. L., Gentles, A. J., Chaudhuri, A. A., Scherer, F., … & Alizadeh, A. A. (2019). Determining cell type abundance and expression from bulk tissues with digital cytometry. Nature biotechnology, 37(7), 773–782.

[106] Han, Y., Wang, Y., Dong, X., Sun, D., Liu, Z., Yue, J., … & Wang, C. (2023). TISCH2: expanded datasets and new tools for single-cell transcriptome analyses of the tumor microenvironment. Nucleic acids research, 51(D1), D1425–D1431.

[107] Leon-Ferre, R. A., Jonas, S. F., Salgado, R., Loi, S., De Jong, V., Carter, J. M., … & International Immuno-Oncology Biomarker Working Group. (2024). Tumor-Infiltrating Lymphocytes in Triple-Negative Breast Cancer. JAMA, 331(13), 1135–1144.

[108] Ali, H. R., Chlon, L., Pharoah, P. D., Markowetz, F., & Caldas, C. (2016). Patterns of immune infiltration in breast cancer and their clinical implications: a gene-expression-based retrospective study. PLoS medicine, 13(12), e1002194.

[109] Chowdhury, A., Pharoah, P. D., & Rueda, O. M. (2023). Evaluation and comparison of different breast cancer prognosis scores based on gene expression data. Breast Cancer Research, 25(1), 17.

[110] Yu, Q., Li, Y. Y., & Chen, Y. (2025). scMalignantFinder distinguishes malignant cells in single-cell and spatial transcriptomics by leveraging cancer signatures. Communications Biology, 8(1), 504.

[111] Patel, A. P., Tirosh, I., Trombetta, J. J., Shalek, A. K., Gillespie, S. M., Wakimoto, H., … & Bernstein, B. E. (2014). Single-cell RNA-seq highlights intratumoral heterogeneity in primary glioblastoma. Science, 344(6190), 1396–1401.

[112] Gao, R., Bai, S., Henderson, Y. C., Lin, Y., Schalck, A., Yan, Y., … & Navin, N. E. (2021). Delineating copy number and clonal substructure in human tumors from single-cell transcriptomes. Nature biotechnology, 39(5), 599–608.

